# Pathogenic KIF1A variants differentially disrupt axonal trafficking and impede synaptic development

**DOI:** 10.64898/2026.01.14.699478

**Authors:** Jayne Aiken, Carris Borland, Nicolas Marotta, Benjamin L. Prosser, Erika L. F. Holzbaur

## Abstract

The nervous system relies on billions of neurons connected through trillions of synapses to support a vast array of vital functions. Despite the critical importance of this synaptic network, the cellular mechanisms dictating synapse formation during human neurodevelopment remain unclear. Long-distance trafficking of synaptic components is critical for both synaptogenesis and the maintenance of synaptic function across lifespan. The microtubule motor KIF1A has a highly conserved role in the trafficking of synaptic vesicle precursors, while mutations in *KIF1A* are causal for the neurodevelopmental and neurodegenerative disease *KIF1A*-Associated Neurological Disorder (KAND). Here, we employ isogenic human induced pluripotent stem cells (iPSCs) gene-edited to express pathogenic *KIF1A* variants to assess how disparate mutations alter synaptic trafficking and function. We compared the effects of both loss-of-function and gain-of-function mutations on KIF1A motor activity. We found that both null (p.C92*) and hypoactive (p.P305L) mutations induce delayed neurite outgrowth, mislocalization of synaptic cargos, and decreased synapse density. Conversely, the hyperactive KIF1A mutation (p.R350G) supports neurite outgrowth but leads to aberrant motility of synaptic vesicle precursors along the axon. Further, live imaging reveals that hyperactive KIF1A induces deficits in the microtubule-dependent patterning of presynaptic components along the developing axon, suggesting a failure to respond to cytoskeletal cues directing cargo delivery. Functional analysis of neuronal activity via multi-electrode arrays reveals delayed synaptic maturation in loss-of-function mutations (p.P305L, p.C92*). In contrast, the hyperactive p.R350G mutation exhibits accelerated activity maturation and possible excitotoxicity. Together, these data provide insights detailing how pathogenic variants in KIF1A causative for KAND exhibit distinct effects at the molecular level that lead to significant downstream deficits in synaptic function in human neurons.

## INTRODUCTION

The human nervous system relies on synaptic communication between neurons to send and receive information. These synaptic connections form intricate neural circuits that underpin all bodily functions including movement, behavior, and thought. Mutations that disrupt synaptogenesis or synaptic function are causal for neurodevelopmental and neurodegenerative diseases including *KIF1A*-Associated Neurological Disorder (KAND). Patients with KAND display a spectrum of symptoms affecting the central and peripheral nervous system, including brain malformation, epilepsy, autism, intellectual disability, and motor disorders (Boyle et al., 2021; Kaur et al., 2020; Klebe et al., 2012, 2006; Nicita et al., 2020). KAND is caused by diverse mutations throughout the *KIF1A* gene, which encodes a member of the kinesin superfamily (Okada et al., 1995).

During development, neurons extend long axonal processes that establish synapses with post-synaptic partners. Microtubules serve as tracks for the active trafficking of synaptic components from sites of synthesis within the soma to presynaptic sites along the axon where their function is required (Aiken and Holzbaur, 2021). Microtubules exhibit intrinsic polarity, with a dynamic plus-end that undergoes bouts of growth and shrinkage (Mitchison and Kirschner, 1984). Axonal microtubules are uniformly oriented plus-end out which helps direct the movement of microtubule motor proteins (Baas et al., 1988; Yogev et al., 2016). Anterograde movement from the soma outward is driven by kinesins that walk toward dynamic microtubule plus-ends. Retrograde movement from the distal axon to the soma is carried out by the motor protein dynein, which walks towards the microtubule minus end. Of the 45 kinesin genes encoded within the human genome, kinesin-3 family member *KIF1A* stands apart due to its essential role in neurodevelopment and its highly motile behavior, which causes it to be colloquially referred to as a ‘marathon runner’ (Soppina et al., 2014). Presynaptic cargos have long been known to rely on KIF1A for delivery (Hall and Hedgecock, 1991; Okada et al., 1995; Otsuka et al., 1991). More recent studies have highlighted the interplay between the regulation of axonal microtubule dynamics and KIF1A-mediated presynaptic delivery (Aiken and Holzbaur, 2024; Guedes-Dias et al., 2019). Microtubules are arranged in a tiled array (Baas et al., 1988) with dynamic plus-ends locally enriched at presynaptic accumulation sites along the developing axon (Aiken and Holzbaur, 2024). Highly processive KIF1A motors traversing axonal microtubules exhibit enhanced detachment frequencies when encountering dynamic plus ends; this enhanced detachment influences the presynaptic accumulation of their cargos (Aiken and Holzbaur, 2024; Guedes-Dias et al., 2019).

Close to 200 distinct disease-causing variants have been identified within *KIF1A*, with patients exhibiting a broad spectrum of clinical phenotypes that roughly correlate to molecular phenotype (Boyle et al., 2021; Sudnawa et al., 2024). Most pathogenic mutations in KIF1A map to the motor domain responsible for binding to microtubules and driving processive movement (Figure 1A). Extensive work including *in vitro* analysis of reconstituted mutant KIF1A motors, studies in model organisms including *C. elegans*, and live imaging in immortalized cell lines have revealed that disease-causing variants within the KIF1A motor domain can impact motility in disparate ways (Boyle et al., 2021; Chiba et al., 2019; Guedes-Dias et al., 2019; Kaur et al., 2020; Shewale et al., 2023). Mutations can cause KIF1A to be hypoactive, traversing microtubules more slowly or less processively, or alternatively induce hyperactivity, leading to faster, more processive movement. Disease-associated mutations to the KIF1A motor domain can also inhibit microtubule binding, or, conversely, promote an aberrant, rigor-bound state where the motor is ‘stuck’ to the microtubule. Less common pathogenic mutations have been identified outside the motor domain, typically in the region immediately preceding or within the Pleckstrin Homology (PH) domain (Boyle et al., 2021; Nicita et al., 2020). The PH domain mediates interactions with membrane lipids (Klopfenstein and Vale, 2004) to facilitate the transport of specialized neuronal organelles carrying synaptic vesicle proteins known as synaptic vesicle precursors (SVPs). Most pathogenic *KIF1A* variants are dominant heterozygous missense mutations occurring *de novo*. While uncommon, recessive homozygous missense mutations within the motor domain, as well as nonsense mutations and frameshift mutations altering the C-terminal, cargo-interacting region of the protein, have been identified (Anazawa et al., 2021; Boyle et al., 2021; Chiba et al., 2019; Klebe et al., 2012; Nicita et al., 2020; Shatarupa et al., 2025). Recessive, inherited variants are of special interest, despite their rarity, as the known homozygous *KIF1A* variants—p.A255V, p.R350W, and p.R350G (explored here)—all hyperactivate motor function. This suggests that hyperactive KIF1A motors are pathogenic when both copies are impacted. Developing an improved understanding of how distinct molecular changes in KIF1A disrupt cellular pathways to cause specific clinical phenotypes will be instrumental in understanding the etiology of KAND. Potentially, these insights will provide guidance in the development of therapeutic strategies that may either be widely useful across KAND patients or instead are tailored for distinct variants.

**Figure 1:**
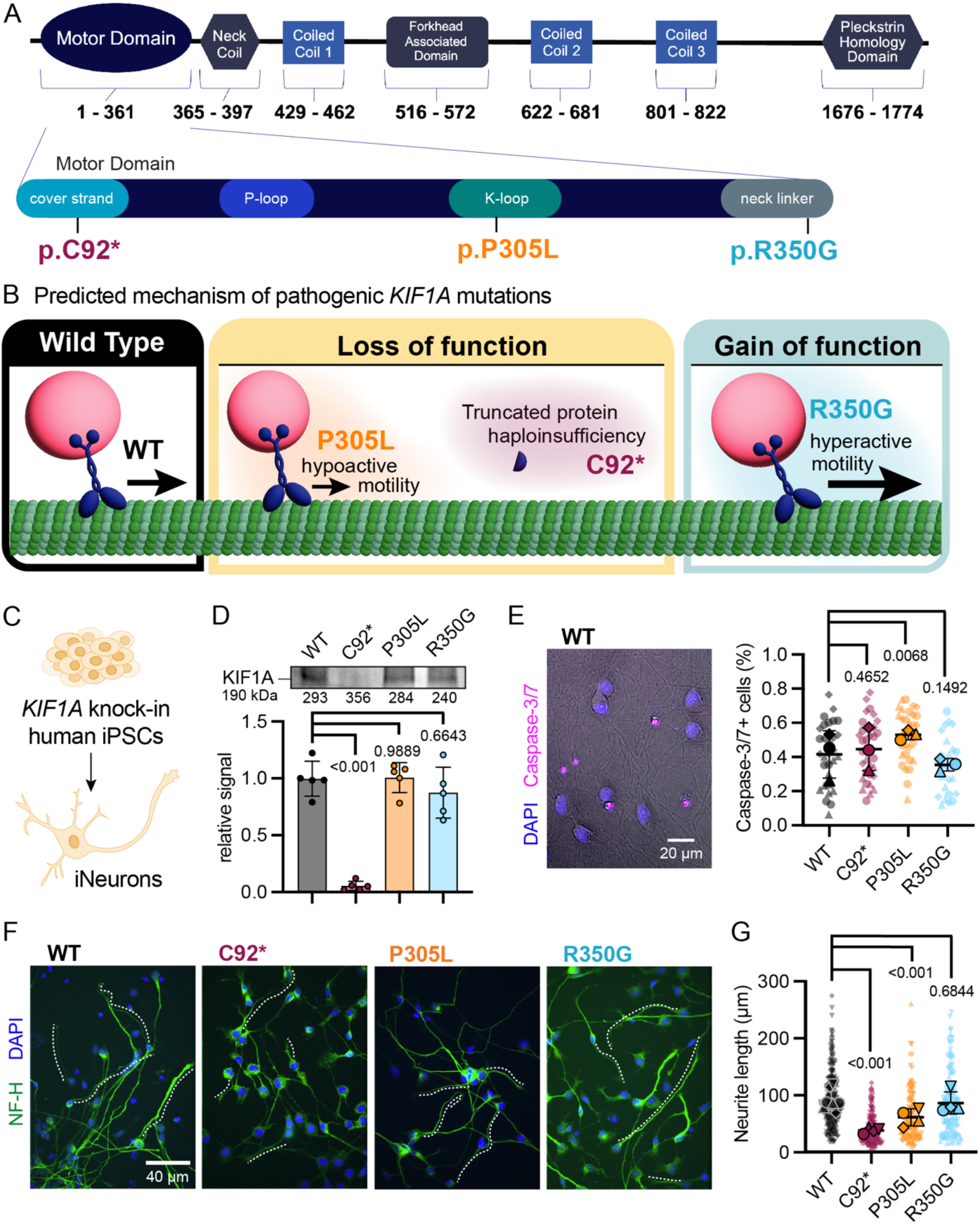
Pathogenic KIF1A mutations impact distinct motor functions and cellular outcomes. **(A)** Mutations to *KIF1A* can lead to a variety of loss- and gain-of-function of disruptions to motor function. Nonsense variant C92* causes early truncation, leading to loss of the mutant allele. Hypoactive variants lead to decreased motor velocity and run length (representative variant P305L). Hyperactive variants cause increase motor velocity and run length (representative variant R350G). Other mutations lead to weakened or enhanced cargo interaction, inability to dimerize or bind to microtubules, or cause the motor to be stuck on the microtubule in a rigor-bound state. **(B)** Annotated schematic of KIF1A protein domains and their corresponding amino acid positions. **(C)** Human iPSCs gene-edited to introduce homozygous p.C92*, p.P305L, or p.R350G to the endogenous *KIF1A* locus of the KOLF2.1J parental line were differentiated into excitatory glutamatergic neurons using tetracycline-inducible expression of NGN2 (iNeurons). **(D)** Example immunoblot and quantification of relative KIF1A protein level in DIV21 wild-type and *KIF1A* mutant iNeurons (mean ± standard deviation; n = 5 independent experiments; reported *p*-values are from ANOVA with multiple comparisons). Total protein levels are provided numerically below the blot. **(E)** Representative image of CellEvent viability assay in DIV7 wild-type neurons (left; Caspase-3/7+ nuclei in magenta, DAPI stained nuclei in blue) and quantification (right) of dead cell fraction for wild-type and *KIF1A* mutant iNeurons (mean ± standard deviation of experimental replicates, n = 30 imaging fields from 3 independent experiments, reported *p*-values determined using linear mixed effect model). **(F, G)** Representative immunocytochemistry images (F) and quantification (G) of neurite extension in wild-type and *KIF1A* mutant iNeurons 24 hours after plating (DAPI nuclear stain, blue; Neurofilament H (NF-H) axon marker, green; mean ± standard deviation of experimental replicates, n =20-30 neurons for each condition from 4 independent experiments, reported *p*-values determined using linear mixed effect model).

Here, we employ isogenic human-induced pluripotent stem cells (iPSCs) gene-edited to endogenously express known pathogenic variants in *KIF1A* and differentiated to cortical-like, glutamatergic neurons (iNeurons; Pantazis et al., 2022) to reveal the cellular consequences of altered KIF1A motor function on synaptogenesis in human neurons. We explore neuronal viability, neurite extension, axonal trafficking, presynaptic cargo distribution, synapse density, and spontaneous synaptic firing capability in iPSC lines that lack KIF1A (p.C92*; Wang et al., 2019), or express hypoactive (p.P305L; Lam et al., 2021; Spagnoli et al., 2019) or hyperactivate (p.R350G; Chiba et al., 2019; Klebe et al., 2012) KIF1A motors. We find that these mutations induce distinct cellular phenotypes. iNeurons lacking *KIF1A* exhibit delayed neurite extension, altered distribution of presynaptic cargos, and fewer synaptic connections compared to isogenic control neurons. iNeurons expressing hypoactive KIF1A motors often phenocopy *KIF1A* null iNeurons, but exhibit decreased neuron viability and significantly altered trafficking of presynaptic cargo. Conversely, hyperactive KIF1A motors support normal neurite outgrowth and viability, but are unable to properly respond to cytoskeletal cues directing stable accumulation of their cargos at presynaptic sites. Despite these mechanistic differences, all three mutations disrupt the appropriate formation of synaptic connections between apposed pre- and postsynaptic compartments, leading to functional changes to synaptic activity. Together, these findings highlight how mutations within the same gene can induce disparate molecular, and cellular, and functional consequences and to provide new insights into the mechanisms by which mutations in the essential neuronal motor *KIF1A* impact synaptic formation and function.

## RESULTS

### Pathogenic *KIF1A* variants cause divergent molecular consequences to motor function

For the last decade variants to the kinesin-3 motor *KIF1A* have been linked to the neurological disorder KAND, with nearly 200 distinct disease-associated mutations now recognized (Boyle et al., 2021; Lee et al., 2015; Nicita et al., 2020; Nieh et al., 2015; Sudnawa et al., 2024). Although predictions can be made based on where a mutation falls within the KIF1A protein, it is difficult to determine the molecular consequence of a variant based on sequence data alone. We therefore turned to recent *in vitro* and *in vivo* functional assays (Boyle et al., 2021; Chiba et al., 2019; Guedes-Dias et al., 2019; Kaur et al., 2020; Shewale et al., 2023) to classify *KIF1A* variants based on molecular phenotype as either loss-of-function or gain-of-function alleles. Loss-of-function may be induced by mutations resulting in hypoactive motility, disrupted cargo binding, disrupted microtubule binding, and rigor binding to microtubules, while gain-of-function alleles induced either hyperactive motility or enhanced binding to cargo.

Based on this analysis, we focused on the in-depth analysis of three lines gene-edited to introduce homozygous mutations into endogenous *KIF1A* locus of the KOLF2.1J parental line by the Cellular Engineering Team at the Jackson Laboratory for Genomic Medicine (JAX). We performed a battery of cellular assays using homozygous *KIF1A* variant lines to more readily assess the potential neurodevelopmental consequences of mutations in this essential neuronal motor. To probe the consequences of KIF1A loss (**Figure 1A, B**), we focused on the nonsense variant p.C92* (Wang et al., 2019), predicted to be a functional null. Variant p.P305L represents the hypoactive motility class of KIF1A mutations; these variants cause decreased motor velocity and run length (Lam et al., 2021; Spagnoli et al., 2019). Variant p.R350G is a hyperactive motor mutation, which leads to increased motor velocity and run length (Chiba et al., 2019; Klebe et al., 2012).

We used tetracycline-inducible expression of *NGN2* to differentiate homozygous, *KIF1A*-edited iPSCs into excitatory glutamatergic neurons, hereafter referred to as iNeurons (**Figure 1C**; Pantazis et al., 2022). Missense mutations p.P305L or p.R350G did not significantly affect levels of KIF1A expression in iNeurons; in contrast, no full length KIF1A protein was detected in p.C92* iNeurons, supporting the prediction that this line is null for KIF1A expression (**Figure 1D**). To support these findings, we also performed RT-qPCR to determine relative mRNA levels for KIF1A using primers recognizing sequences upstream of the p.C92 nonsense mutation (**Figure S1A**). While the p.C92* variant induced the greatest decrease in KIF1A mRNA expression, consistent with nonsense-mediated decay, reduced mRNA levels were observed in all mutant lines (p.C92*, 15% of wild-type; p.P305L, 48%; p.R350G, 36%). The decrease in KIF1A mRNA observed in both the p.P305L and p.R350G lines was surprising given the normal levels of KIF1A protein expressed (**Figure 1D**), but suggests that levels of *KIF1A* mRNA are not the sole determinant for protein expression.

To assess the possibility that expression of another member of the kinesin-3 family of motors might be upregulated in response to decreased levels or functional inhibition of KIF1A , we also determined relative mRNA expression levels of kinesin-3 family members *KIF1B* and *KIF1C*. Unexpectedly, we found that iNeurons homozygous for p.C92* also exhibited decreased mRNA expression of other motors; we observed a significant reduction in KIF1B mRNA and a trend toward a decrease in KIF1C mRNA (**Figure S1A**). Effects on protein expression levels could not be assessed due to inadequate specificity of available KIF1B and KIF1C antibodies, so we focused on differences in relative transcript levels. Variant p.P305L, but not p.R350G, exhibited significantly decreased levels of KIF1B mRNA by qPCR (**Figure S1A**), but no significant changes in expression of either KIF1B or KIF1C were observed for iNeurons expressing the R350G mutation. Together, these data suggest that deficits in KIF1A function do not induce compensatory enhancements in the transcription of the related motors KIF1B and KIF1C.

### *KIF1A* variants differentially affect early stages of neuronal differentiation

Previous studies in mouse models demonstrated loss of *KIF1A*-deficient primary neurons, suggesting KIF1A protein function is essential for neuron survival (Yonekawa et al., 1998). Surprisingly, we found that p.C92* human iNeurons lacking *KIF1A* do not exhibit a significant defect in neuronal viability compared to isogenic controls, as determined at DIV7 with CellEvent Caspase-3/7 detection assay (**Figure 1E**), nor did we see increased cell death in neurons expressing the hyperactive p.350G mutation. In contrast, a significant increase in cell death was observed for neurons expressing the p.P305L allele, suggesting that this hypoactive mutation may function as a dominant negative in this context. These data reveal that KIF1A is not required for the viability of human iNeurons *in vitro*, while expression of hypoactive, but not hyperactive, KIF1A motor activity is detrimental to cell viability.

Neuron polarization and neurite outgrowth are crucial steps in neuronal morphogenesis. To examine the effects of KIF1A mutations on the ability of human neurons to extend nascent neurites, we stained iNeurons 24 hours after plating with antibodies to Neurofilament-H (**Figure 1F**). Neurofilaments are key structural components in neurons that are particularly abundant in axonal projections. We found that p.C92* neurons lacking KIF1A exhibit a striking delay in early neurite extension (**Figure 1G**). The hypoactive p.P305L variant likewise caused a decrease in neurite length 24 hours after plating, though not as severe as observed in p.C92* neurons. In contrast, neurite outgrowth in neurons expressing the hyperactive p.350G mutant was not significantly different than isogenic control neurons. These results suggests that KIF1A is an important regulator of early neurite outgrowth. Thus, while mutations in KIF1A may induce delayed axonal outgrowth in vivo, which can have dramatic consequences on neurodevelopment as axons encounter and respond to time-dependent cues as they traverse the developing nervous system (Zang et al., 2021), either KIF1A loss- or gain-of-function mutations are unlikely to block outgrowth completely.

### Human neurons lacking *KIF1A* can compensate to rescue presynaptic cargo movement

Classic studies in *C. elegans* and mice have identified the principal axonal cargo of KIF1A motors as presynaptic components required for synaptic vesicle formation (Hall and Hedgecock, 1991; Okada et al., 1995; Otsuka et al., 1991; Yonekawa et al., 1998). We therefore investigated how pathogenic mutations in KIF1A impact the axonal transport of SVPs in human neurons by tracking movement of synaptophysin (SYP), a key transmembrane protein in synaptic vesicles (Maeder et al., 2014). We imaged wild-type and *KIF1A* mutant iNeurons expressing the fluorescent reporter mScarlet-SYP at DIV21 to track axonal SVP movement (**Figure 2A, B**). To facilitate visualization of motile SVPs through the axon, we photobleached axonal regions of interest prior to imaging to deplete accumulated mScarlet-SYP signal (Dou et al., 2024; Guedes-Dias et al., 2019).

**Figure 2:**
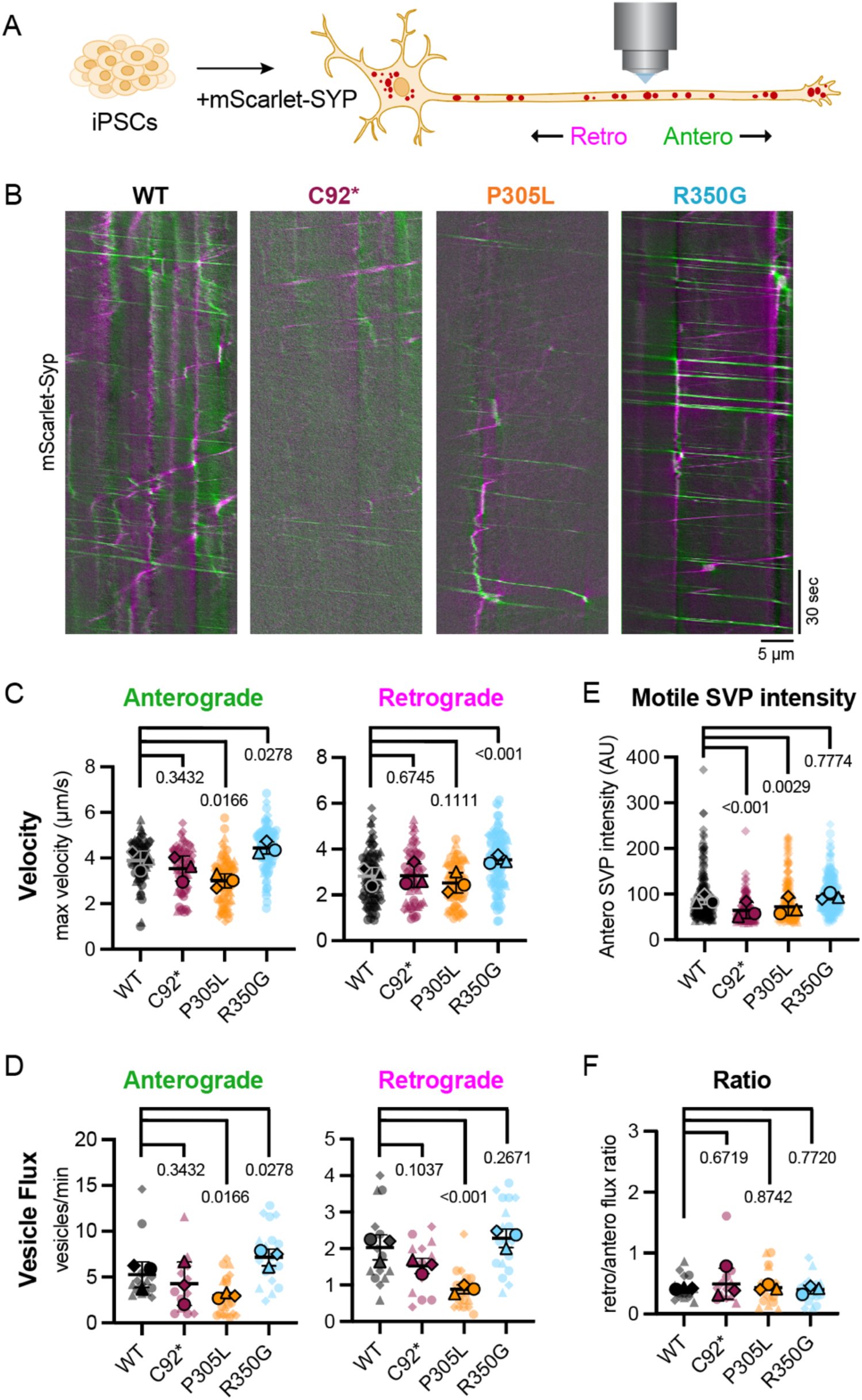
KIF1A mutations alter trafficking behavior of synaptic cargos along developing axons. **(A)** iNeurons transfected with mScarlet-Synaptophysin (Syp) were live-imaged at five frames per second to detect rapid synaptic vesicle precursor (SVP) movement along the axon. Cytoplasmic mScarlet-SYP signal was photobleached prior to live imaging to promote visualization of incoming SYP cargo. **(B)** Representative kymographs of Syp movement in DIV21 wild-type and *KIF1A* mutant iNeurons. Anterograde and retrograde tracks are pseudo-colored green and magenta, respectively. **(C)** Velocity of the five fastest moving SVPs per axon (maximum velocity, µm/s) in the anterograde (left) and retrograde (right) direction in wild-type, C92*, P305L, and R350G iNeurons. **(D)** Mean intensity of motile SVPs moving in the anterograde direction. All motile anterograde SVPs (movement of >10 µm in the anterograde direction) were included in this analysis. **(E)** Vesicle flux (vesicles/minute) for anterograde (left) and retrograde (right) moving SVPs in wild-type and *KIF1A* mutant iNeurons. **(F)** Ratio of retrograde/anterograde flux for each genotype. All plots display mean ± standard deviation of experimental replicates, n = 74 motile SVPs from 3 independent experiments, reported *p*-values determined using linear mixed effect model.

Consistent with previous reports in mouse, rat and human neurons (Aiken and Holzbaur, 2024; De Pace et al., 2020; Dou et al., 2024; Guedes-Dias et al., 2019; Niwa et al., 2008; Rizalar et al., 2023), we observed rapid, highly-processive SVP movement in wild-type iNeuron axons, with an average of five SVPs per minute moving at an instantaneous velocity of ∼4 µm/s. Unexpectedly, we discovered that p.C92* mutant neurons do not exhibit significant changes in the maximum velocity achieved by motile SVPs (**Figure 2C**) or to SVP flux, the number of SVP cargos moving through an axon per minute (**Figure 2D**). These data support the notion that in the absence of *KIF1A*, human neurons rely on a motor with similar motile parameters to compensate for axonal SVP transport. The main contender for this compensatory motor is kinesin-3 family member KIF1Bβ, which was previously shown to traffic SVP cargo (Niwa et al., 2008; Zhao et al., 2001). Given the collective decrease in kinesin-3 mRNA observed in p.C92* iNeurons, encompassing KIF1A, KIF1B, and KIF1C **(Figure S1A)**, this suggests that kinesin-3 mRNA levels are not limiting for SVP axonal transport. While we did not note a decrease in the number of motile SVPs in p.C92* axons, we did find that the fluorescent intensity of motile SVPs in this line was significantly lower than isogenic controls (**Figure 2E**). Therefore, while the number and velocity of SVPs was unaltered, cells lacking KIF1A may exhibit a deficit in the loading and/or the transport of SVPs of appropriate size.

### Hypo- and hyperactive KIF1A variants impact both anterograde and retrograde trafficking of SVPs

Both hypoactive and hyperactive KIF1A mutant motors impact SVP axonal transport behavior, albeit with opposing effects. Supporting previously published data collected using *in vitro* reconstituted systems or in model organisms (Boyle et al., 2021; Kaur et al., 2020; Lam et al., 2021), human iNeurons expressing endogenous p.P305L mutant motors exhibited fewer SVP vesicles moving at slower velocities along the axon (**Figure 2C, D**). Neurons expressing the hypoactive p.P305L mutation exhibited a small yet significant decrease in anterograde velocities, with a trend toward slower retrograde transport. SVP flux was slightly decreased in both directions. As with the p.C92* mutation, the p.P305L mutation induced a decrease in the intensity of motile SVP cargos.

Conversely, the hyperactive motor p.R350G induced an increase in both the velocity and flux of SVPs moving along the axon (**Figure 2C, D**). Perturbations in axonal transport were observed for the anterograde SVP population, which relies on kinesin-based motility, as well as the dynein-driven, retrograde SVP population. Strikingly, the p.R350G hyperactive mutant caused a dramatic increase in maximum retrograde velocity (increased 3.5 µm/s compared to 2.8 µm/s in wild-type). These surprising results suggest that changes to the anterograde motor KIF1A impact the motile behavior of dynein, responsible for retrograde movement. Changes in SVP flux were observed in the anterograde and retrograde directions for both the p.P305L and p.R350G variants. For variant p.P305L, decreased flux was observed in both the anterograde and retrograde directions, while p.R350G variant induced increased SVP flux in both the anterograde and retrograde directions. In all cases, the ratio of incoming vs. outgoing SVPs remained unchanged across genotypes (**Figure 2F**). This suggests that the amount of outgoing cargo carried by KIF1A directly influences the amount of incoming cargo carried by dynein.

### *KIF1A* variants significantly alter the compartmental distribution of presynaptic cargos in human iNeurons

Next, we set out to determine the ramifications of altered SVP transport on cargo distribution throughout the neuron. We employed immunoblotting and immunocytochemistry to detect endogenous presynaptic proteins synaptophysin (SYP) and synaptobrevin-2 (SYB2, also known as VAMP2) and their relative distribution to either the soma or presynaptic sites, modeled here using heterologous presynapses generated in response axonal contacts with neuroligin-1-expressing HEK293 cells (Aiken and Holzbaur, 2024; Biederer and Scheiffele, 2007; Dou et al., 2024). This heterologous presynapse assay was used to determine how *KIF1A* variants impact the accumulation of presynaptic components along the axon, to examine the spatiotemporal control of presynaptic formation (Aiken and Holzbaur , 2024). Overall protein expression levels of SYP and SYB2 remain unchanged across conditions (**Figure 3A**, **Figure S1B**). However, the distribution of these presynaptic proteins is significantly altered in neurons expressing *KIF1A* mutants. Specifically, iNeurons expressing loss-of-function variants p.C92* (null) and p.P305L (hypoactive motor) exhibit aberrant retention of synaptic vesicle cargos in the soma (**Figure 3B-E**). In parallel, these iNeurons exhibit significant depletion of these key synaptic components from heterologous presynapses (**Figure F-I**). In contrast, expression of the hyperactive p.R350G motor mutation induced to the opposite distribution, with lower levels of SYB2 and SYP in the soma (**Figure 3B-E**)and significantly increased levels of SYP accumulation at heterologous presynaptic sites (**Figure F-I**).

**Figure 3:**
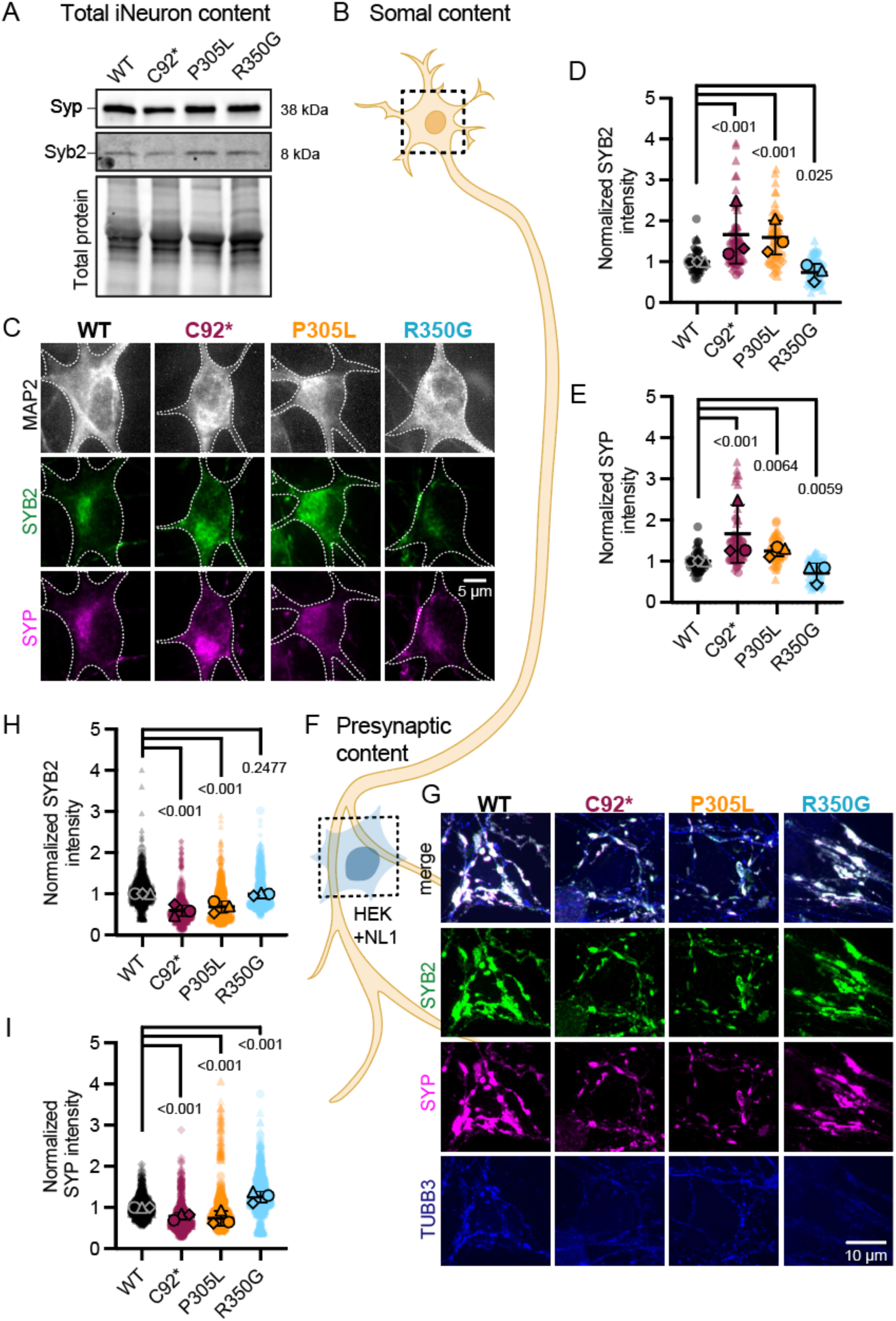
Loss-of-function vs. gain-of-function KIF1A mutations lead to opposite synaptic cargo distribution defects. **(A)** Representative immunoblot of endogenous synaptophysin (Syp) and synaptobrevin-2 (Syb2) protein in wild-type and *KIF1A* mutant iNeurons at DIV21. No significant change is observed for total neuronal protein levels for either synaptic cargo (**Figure S1B**). **(B-E)** Examination of subcellular Syp and Syb2 content in the iNeuron soma (A): Representative immunocytochemistry images (B) of somatodendritic compartment (MAP2, white) and synaptic cargos (SYB2, green; SYB2, magenta). Quantification of mean soma intensity normalized to wild-type for SYB2 (D) and SYP (E). Plots display mean ± standard deviation of experimental replicates, n = 30 neurons from 3 independent experiments, reported *p*-values determined using linear mixed effect model. **(F-H)** Examination of subcellular Syp and Syb2 content at heterologous presynaptic compartments formed in response to neuroligin-1-expressing HEK cells (F): Representative immunocytochemistry images (G) of heterologous presynapses formed on NL1+ HEK cells (SYB2, green; SYP, magenta; TUBB3 axonal marker, blue) in wild-type and *KIF1A* mutant iNeurons. Quantification of mean soma intensity normalized to wild-type for SYB2 (D) and SYP (E). Plots display mean ± standard deviation of experimental replicates, n = 30 HEK cells interacting with numerous axons from 3 independent experiments, reported *p*-values determined using linear mixed effect model.

### *KIF1A* variants disrupt accumulation of presynaptic cargo along the axon

KIF1A-mediated presynaptic cargo transport along the axon is regulated by the periodic enrichment of dynamic microtubule ends that promote the local delivery of SVPs (Aiken and Holzbaur, 2024; Guedes-Dias et al., 2019). Due to its highly processive motility, KIF1A motors typically traverse the microtubule polymer processively toward dynamic plus ends, where they preferentially detach due to a lower affinity for GTP-bound tubulin dimers enriched within the lattice at these sites (Guedes-Dias et al., 2019). The distribution of microtubule ends along the axon therefore act as delivery cues for incoming SVPs carried by KIF1A (Guedes-Dias et al., 2019; Aiken and Holzbaur, 2024). We hypothesized that mutations that disrupt KIF1A’s motile properties would impact the motor’s ability to respond to appropriate cues for SVP distribution.

To test this hypothesis, we live imaged DIV21 wild-type and *KIF1A* mutant iNeurons expressing a bicistronic vector that simultaneously expresses microtubule plus-end marker GFP-MACF43 and SVP protein mScarlet-SYB2 (**Figure 4A**). MACF43 binds to and tracks growing microtubule ends, facilitating the visualization of microtubule ‘comets’ (Yau et al., 2016), while SYB2 allows us to visualize both motile SVPs and those that accumulate at protosynaptic sites along the axon (Aiken and Holzbaur, 2024). We acquired images at 1 frame per second to visualize both the GFP and mScarlet channels. This acquisition rate is sufficiently fast to detect the growth of dynamic microtubule ends and to visualize static accumulations of SVPs along the axon, but not to monitor the very rapid movement of motile SVPs shown in **Figure 2**. Comparisons across genotypes indicate that the plus-end dynamics of axonal microtubule are not significantly altered in *KIF1A* mutant iNeurons, as we observed no significant differences across the cell lines tested (**Figure 4B, C**). Consistent with our previous reports in primary rat neurons (Guedes-Dias et al., 2019) and in human neurons generated from a distinct iPSC parental line (WTC11; Aiken and Holzbaur, 2024), microtubule comets within the axon are typically enriched at sites coincident with SVP accumulation, with cycles of recurrent microtubule plus-end growth detected over time.

**Figure 4:**
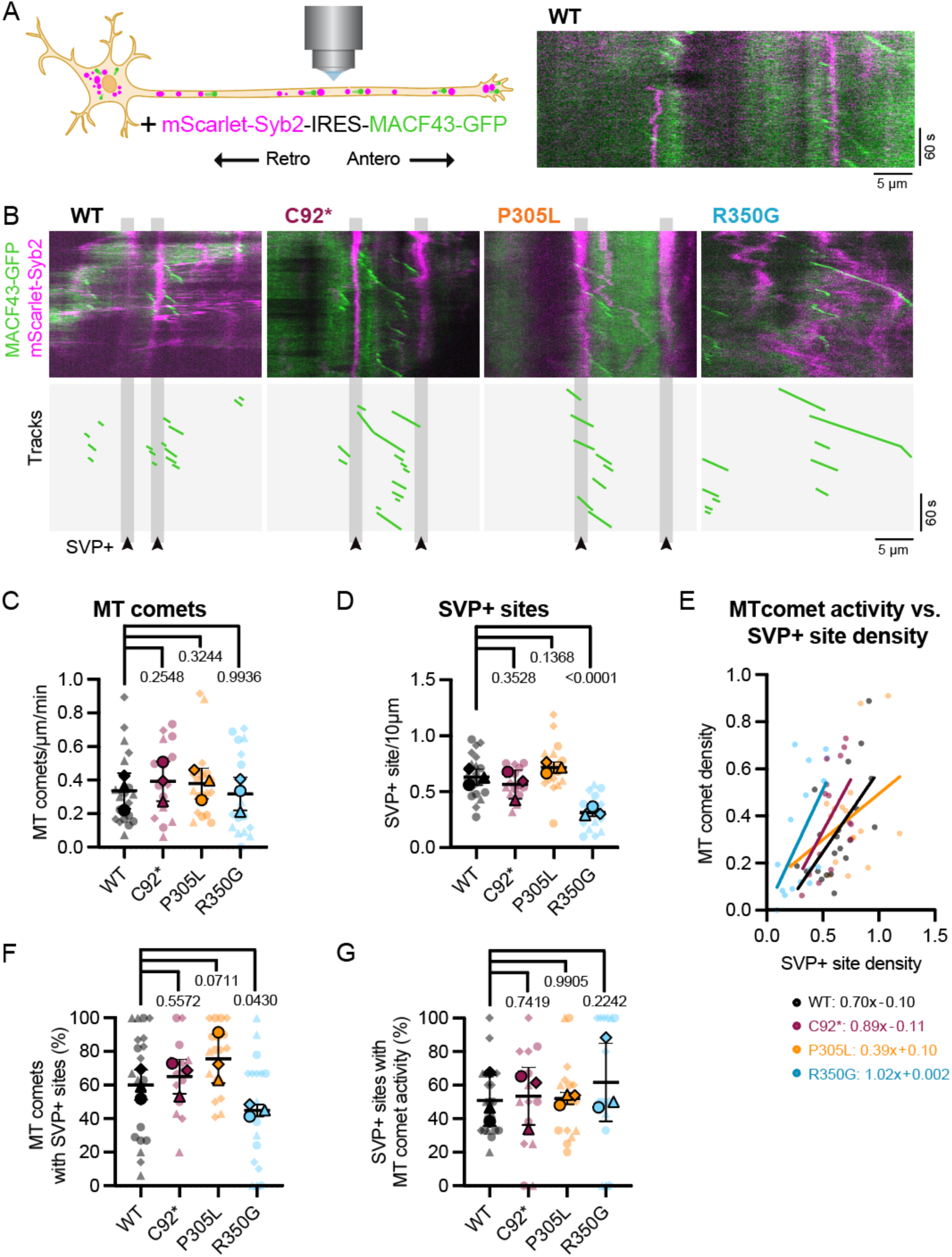
Hyperactive KIF1A mutant disrupts axonal presynaptic cargo delivery despite appropriate microtubule cues. **(A)** iNeurons at DIV21 expressing bicistronic vector mScarlet-Syb-IRES-MACF43-GFP were live-imaged at one frame per second to detect SVP accumulations and microtubule polymerization along the axon. Right panel: example wild-type kymograph containing two regions of recurrent microtubule dynamics associated with the accumulation of stable SVPs. **(B)** Representative kymographs (top panels) and tracks (lower panels) of SVP+ sites and microtubule comet events in wild-type and *KIF1A* mutant iNeurons. **(C)** Microtubule comet activity (MT comet/µm/min) in wild-type and *KIF1A* mutant axons. No significant difference in frequency of microtubule growth is observed across genotype. **(D)** Presynaptic accumulation site density (SVP+ sites/10 µm) in wild-type and *KIF1A* mutant axons. Hyperactive mutant R350G leads to significantly fewer stable SVP+ sites within the axon. **(E)** Microtubule comet activity plotted against SVP+ site density for wild-type and *KIF1A* mutant axons. Each point within the scatterplot represents a distinct axon, with regression lines displayed and equations provided below. R^2^ values: wild-type=0.38; C92*=0.30; P305L=0.16; R350G=0.21. **(F)** Percentage of total microtubule comets associated with presynaptic accumulations (comets that initiate, terminate, or pass through SVP+ sites). **(G)** Percentage of presynaptic accumulations that experience microtubule comets activity. All plots display mean ± standard deviation of experimental replicates, n = 15 axons from 3 independent experiments, reported *p*-values determined using linear mixed effect model.

While microtubule dynamics were not significantly altered across genotypes, we observed a striking change in the density of accumulated synaptic vesicle precursor sites (SVP+ sites) in axons expressing the within hyperactive *KIF1A* p.R350G allele (**Figure 4D**). While *KIF1A* p.R350G iNeurons typically express high levels of synaptic markers and exhibit robust trafficking of SVPs along the axon (**Figure 2C-E**, **Figure 3H-I**, **Figure 4B**), we find that SVPs fail to accumulate at discrete positions along the axon. Both isogenic control and p.C92* *KIF1A* null axons exhibit ∼0.6 stable SVP+ site per 10 µm, while in neurons expressing the hyperactive *KIF1A* p.R350G mutant this density is decreased by 50% to 0.31 sites/10 µm (**Figure 1D**). The opposite finding was observed for neurons expressing the hypoactive p.P305L mutant, which exhibited a trend toward an increased number of stable SVP+ sites (0.72 sites/10 µm).

When SVP+ site density is plotted against microtubule comet activity, a general linear relationship is observed (**Figure 4E**; Aiken and Holzbaur, 2024). This relationship stems from the observation that SVP+ sites house most of the axonal microtubule comet activity (**Figure 4F**), with stretches of axons lacking SVP+ sites containing few dynamically growing microtubules and SVP+-enriched axonal regions exhibiting numerous recurrent microtubule comet events. Disrupting KIF1A motility perturbs this relationship. Expression of the hyperactive p.R350G mutant leads to a leftward shift in the linear regression line, representing the decreased SVP+ sites despite normal microtubule comet activity. Both hyperactive and hypoactive *KIF1A* mutations decreased the R^2^ value for the line of best fit, disrupting the general relationship between microtubule plus-end growth and SVP accumulation (R^2^ values: wild-type=0.38; C92*=0.30; P305L=0.16; R350G=0.21). Due to the reduction of SVP+ site formation in p.R350G axons, significantly more microtubule comet events are found outside of these stable presynaptic accumulations (**Figure 4F**). However, when SVP+ sites are formed in hyperactive *KIF1A* p.R350G axons, they are as likely to house recurrent microtubule comet events as wild-type SVP+ sites (**Figure 4G**). These data underscore the importance of appropriate motor response to cytoskeletal cues during axon development for presynaptic cargo distribution. As neuronal arbors mature, this pre-patterning of synaptic cargo unloading zones may functionally direct future synapse biogenesis and form the basis for continued synaptic function.

### Pathogenic *KIF1A* mutations disrupt synaptic biogenesis

Next, we asked how *KIF1A* mutations might alter presynaptic distribution. To promote *in vitro* synapse formation, we co-cultured human iNeurons with primary glia enriched for astrocytes (Schildge et al., 2013) and allowed the cultures to mature to DIV42. Pre- and postsynaptic puncta were detected through immunocytochemistry (**Figure 5A**); we looked for apposition of the presynaptic marker Synapsin I/II (SYN, magenta), the post-synaptic marker PSD-95 (green), and the somatodendritic compartment (MAP2; blue).

**Figure 5:**
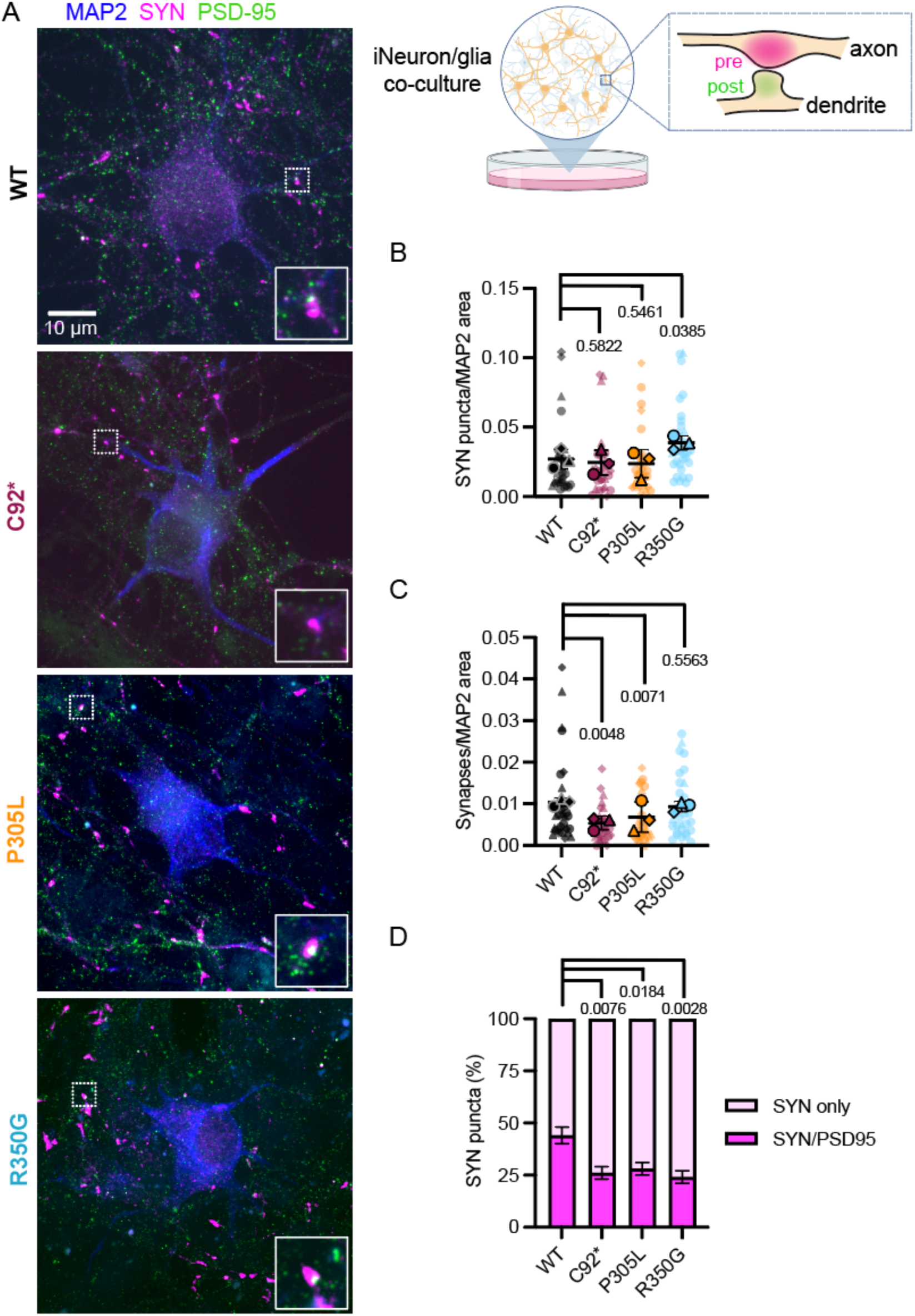
Pathogenic KIF1A mutations disrupt the formation of *bona fide* synapses consisting of pre- and post-synaptic compartments. **(A)** Experimental schematic and representative immunocytochemistry images of presynaptic synapsin I/II (SYN, magenta), postsynaptic (PSD-95, green), and somatodendritic compartments (MAP2, blue) in wild-type, *KIF1A-C92\**, -*P305L*, and -*R350G* iNeurons co-cultured with primary rat astrocytes. Insets display representative presynaptic accumulations apposed to postsynaptic puncta (WT, P305L, and R350G) or alone (C92*). **(B, C)** Syn puncta density (C) and *bona fide* synapse density consisting of apposed pre/postsynaptic puncta (D) normalized to MAP2 area for wild-type or *KIF1A* mutant iNeurons. Pre- and postsynaptic puncta were defined using SynapseJ quantification on z-stack images. Plots display mean ± standard deviation of experimental replicates, n = 30 imaging fields from 3 independent experiments, reported *p*-values determined using linear mixed effect model. **(D)** Percentage of SYN puncta apposed to a postsynaptic compartment (magenta) or alone (light pink). Despite differences observed for SYN puncta/synapse density between loss-of-function and gain-of-function *KIF1A* mutants, each mutation disrupts the ratio of presynaptic puncta aligning with postsynaptic partners. Reported *p*-values were determined using Chi-Square test of independence.

Both the p.C92* and p.P305L hypoactive mutations in KIF1A led to fewer *bona fide* synaptic connections, defined as apposed pre- and post-synaptic compartments. However, the overall number of SYN-positive puncta was not altered, consistent with our live imaging analysis of SVP+ sites (**Figure 4**). The expression of hyperactive KIF1A p.R250G motors led to an increase in SYN-positive presynaptic puncta, but no change in apposed pre- and postsynaptic partners (**Figure 5B, C**). Overall, we observed synapse formation at roughly similar rates across *KIF1A* alleles tested, although we did see a consistent decrease in the ratio of presynaptic puncta apposed to post-synaptic markers in the pathogenic *KIF1A* variants compared to our isogenic control line (**Figure 5D**). This points to a mild deficit in synaptic formation upon disruption of KIF1A function, with potentially a longer-term deficit in synaptic maintenance. This led us to screen for potential deficits in synaptic function.

### Pathogenic *KIF1A* variants induce functional synaptic firing defects in human neurons

To assess the functional impact of pathogenic KIF1A motors on the synaptic function of human neurons, we employed multi-electrode array (MEA) electrophysiology. MEA technology provides a powerful system for studying neurodevelopment and synaptogenesis due to its ability to longitudinally monitor neuronal activity of neuronal cultures. Applying this approach to iNeurons in monoculture, we found that loss-of-function *KIF1A* alleles p.C92* and p.P305L exhibited a decrease in synaptic firing, while no change was observed in spontaneous activity for the hyperactive p.R350G variant (**Figure 6A, B**).

**Figure 6:**
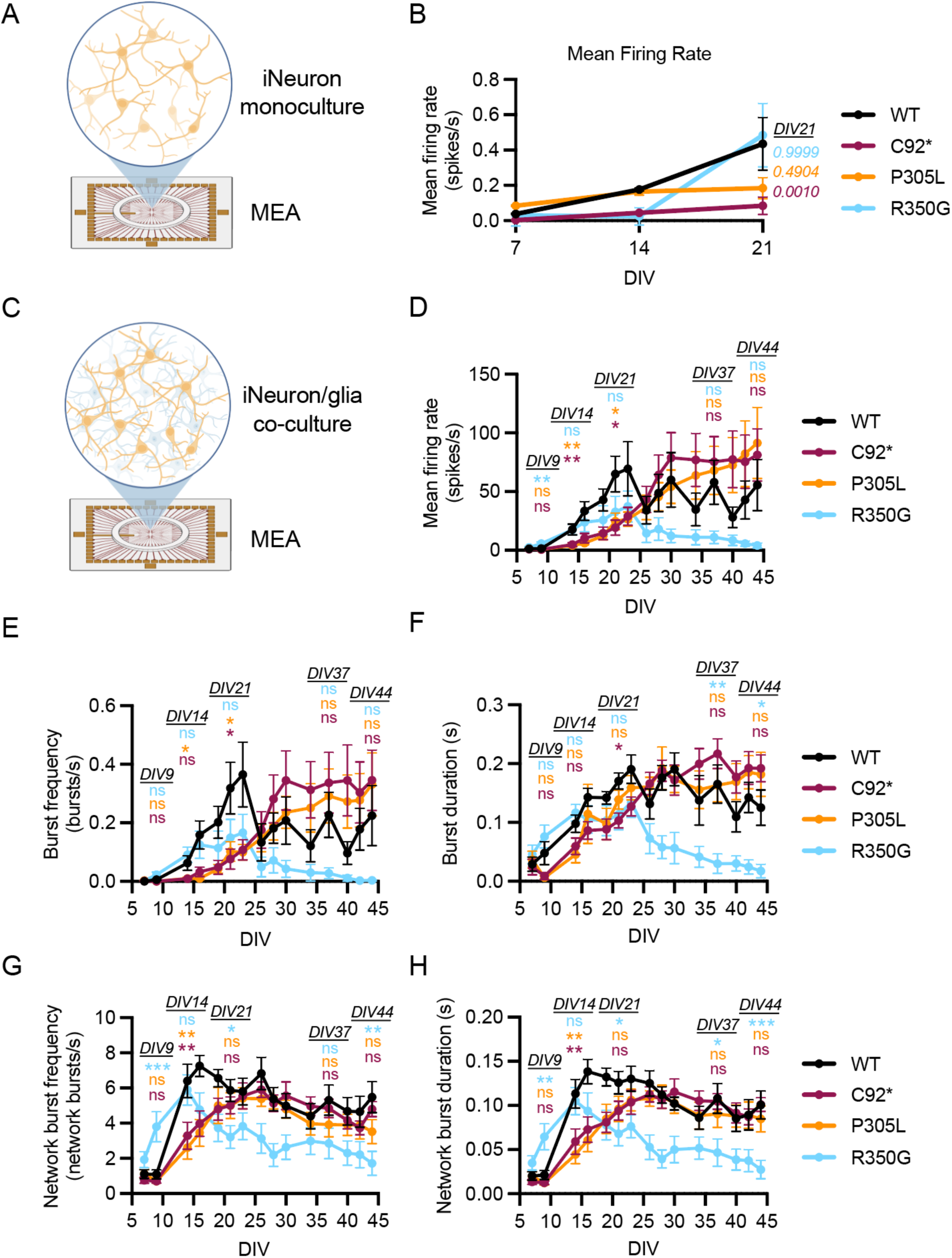
*KIF1A* loss-of-function vs. gain-of-function variants lead to differing developmental impacts on neuronal activity. **(A)** Schematic of synaptic recording experimental set-up. Wild-type and *KIF1A* mutant iNeurons were cultured on multi-electrode arrays (MEA) and recorded weekly to observe spontaneous firing rate. **(B)** Longitudinal tracking of mean firing rate (spikes/second) for wild-type and *KIF1A* mutant iNeurons. Wild-type and hyperactive *KIF1A* mutant iNeurons grown in monoculture exhibit increasing, albeit modest, spontaneous synaptic release from DIV7-21. Kruskal-Wallis nonparametric test followed by Dunn’s test for multiple comparisons was performed to compare DIV21 mean firing rates, with p-values provided. **(C)** Schematic of synaptic function experimental set-up for wild-type and *KIF1A* mutant iNeurons co-cultured with rat astrocytes for MEA recording. Spontaneous firing rate was recorded longitudinally as iNeurons mature. **(D)** Wild-type and KIF1A mutant iNeurons co-cultured with rat astrocytes exhibit increased mean firing rate (spikes/second) over developmental time. WT cultures exhibited an increase in spikes over longitudinal recordings, plateauing around DIV24. KIF1A pathogenic variants impacted the developmental trend in mean firing rate. **(E, F)** Burst frequency (E) and duration (F) for WT, p.C92*, p.P305L, and p.R350G conditions support the conclusion that loss-of-function variants delay synaptic maturation and the hyperactive variant. **(G, H)** Network Burst frequency (G) and duration (H) recorded longitudinally reveals precocious activity in p.R350G iNeurons and delayed maturation in p.C92* and p.P305L. Synaptic bursts are defined as repeated spikes on an electrode that occur within 50ms and terminate after 100ms of inactivity, and network bursts counted as simultaneous bursts detected across at least 4 electrodes. Statistical analyses were conducted for developmental timepoints across the experiment, as denoted in the figure. Note: one WT replicate was lost after DIV23, affecting panels D-H. P-values are indicated based on a One-way ANOVA with Dunnett’s multiple comparison test: * = 0.05; ** = 0.01, *** = 0.0001, **** <0.0001. For full distribution of data and specific p-values at these timepoints, see Figure S2.

**Figure 7.**
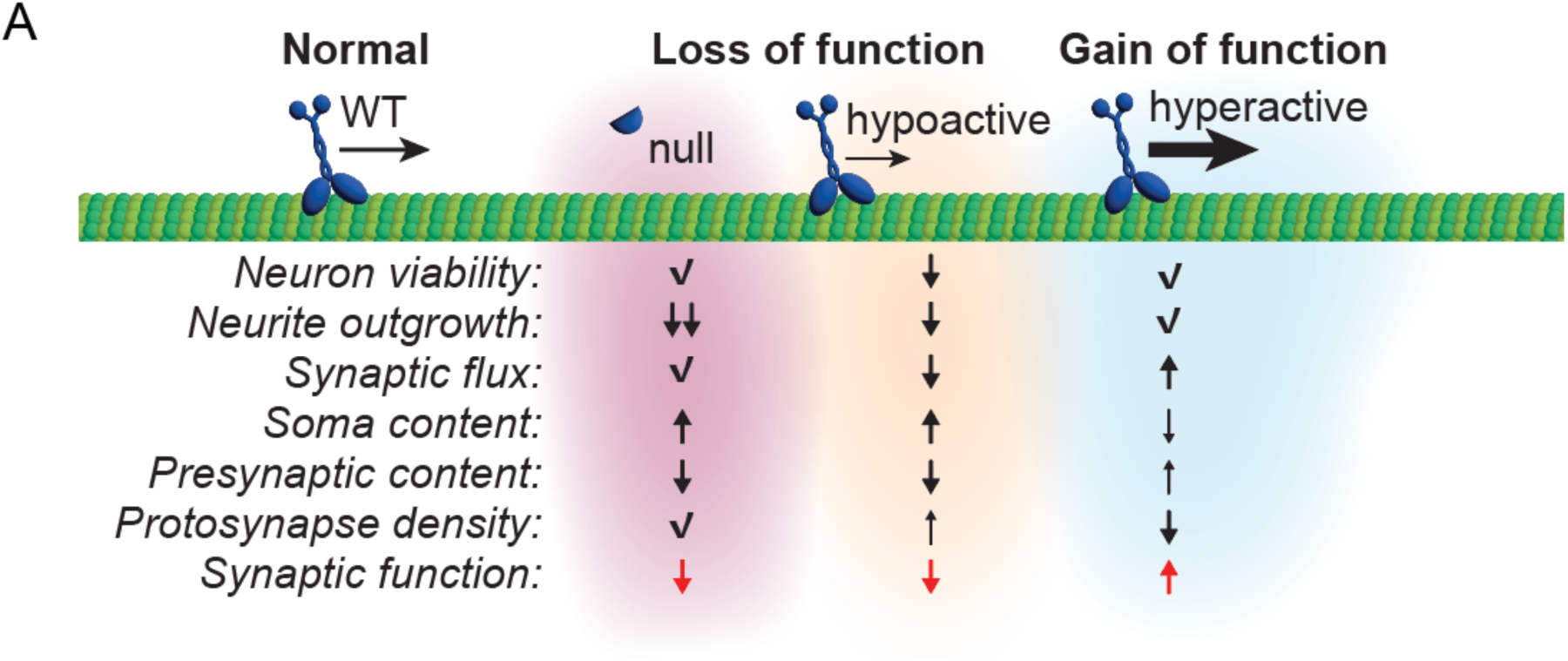
(A) Summary: Distinct changes to KIF1A function differentially impact cellular phenotypes to lead to synaptic activity defects.

However, recent work has shown that iNeurons develop more robust synapses when co-cultured with primary astrocytes (Aiken and Holzbaur, 2024; Benner et al., 2025; Meijer et al., 2019; **Figure 6C**). In our co-culture MEA experiments we monitored multiple parameters of synaptic activity including mean firing rate (**Figure 6D**), burst frequency and duration (**Figure 6E, F**), and network burst frequency and duration (**Figure G, H**). Overall, we noted that co-culturing iNeurons with astrocytes led to a 100-fold increase in spontaneously recorded spikes per second, emphasizing the previously reported importance of astrocytes in promoting synapse formation and function in human neurons (Aiken and Holzbaur, 2024; Faissner et al., 2010; Hedegaard et al., 2020; Liu et al., 2021).

Both of the loss-of-function alleles (p.C92* and p.P305L) exhibited a delay in synaptic maturation compared to WT cultures. While WT iNeurons consistently exhibit a gradual increase in mean firing rate, bursts, and network bursts over time until reaching a plateau around DIV23, neurons expressing the p.C92* and p.P305L mutations exhibited a more prolonged maturation, eventually reaching peak activity around DIV28. In contrast, the hyperactive allele p.R350G initially induced precocious network burst activity indicative of accelerated maturation, reaching a peak around DIV13-15. Following this peak in activity, however, p.R350G neurons demonstrated a sharp decline in activity evident by DIV20, potentially due to excitotoxicity. Together, these data demonstrate that pathogenic variants in *KIF1A* result in distinct cellular perturbations that differentially affect synaptic activity across developmental time. From a functional standpoint, these observations indicate that tuning KIF1A motor activity in either direction, towards either hypo- or hyper-activity, detrimentally impacts synaptic development in human neurons.

## DISCUSSION

A growing number of pathogenic variants have been identified within the KIF1A gene, linked to a broad spectrum of developmental and degenerative symptoms collectively referred to as KAND (Boyle et al., 2021). More recently, KIF1A mutations have also been identified in patients with ALS and atypical parkinsonism (Bernard et al., 2024; Homayoun et al., 2025). The majority of these pathogenic variants are heterozygous, de novo missense mutations (Boyle et al., 2021). *In vitro* and *in vivo* studies implicate both loss-of-function and dominant negative mechanisms can lead to pathogenesis (Boyle et al., 2021; Rao et al., 2025).

Accumulating biophysical data indicate that mutations in KIF1A have the capacity to disrupt motor function through discrete molecular mechanisms, affecting microtubule binding, cargo binding, motor velocity, and motor processivity (Nieh et al., 2015; Chiba et al., 2019; Boyle et al., 2021; Lam et al., 2021; Rao et al., 2015). The impact of KIF1A mutations have also been tested in model organisms, including *C. elegans* and iPSC-derived motor neurons, identifying deficits in axonal transport, protein aggregation, and autophagy (Anazawa et al., 2022; Chiba et al., 2019; Zhao et al., 2024). However, a careful dissection of the consequences of KIF1A mutations on human synaptic development and function at the cellular level has not yet been undertaken. Here, we directly compared three isogenic human iPSC lines gene-edited to express KIF1A mutations with increasing levels of predicted severity. We differentiated these neurons to glutamatergic cortical-like neurons and compared the effects of a null mutation (p.C92*), a mutation that induces hypoactive motor function (p.P305L), and a mutation leading to hyperactive motor function (p.R350G). We find that these variants induce distinct effects on neuronal viability, neurite outgrowth, SVP trafficking, synaptic density, and synaptic activity deficits. These findings are in line with the broad and diverse spectrum of disease observed in KAND patients, who present with a range of neurologic features and clinical severity. These data suggest that patients will require personalized therapeutic strategies, as distinct pathogenic variants differentially disrupt synaptic development across molecular, cellular, and functional scales.

### Pathogenic mutations in KIF1A have disparate effects on motor function

Almost 200 pathogenic mutations in KIF1A have been identified to date (Sudnawa et al., 2024). While mutations have been identified throughout the protein, pathogenic variants tend to cluster within the motor domain (Boyle et al., 2021; Sudnawa et al., 2024). Careful comparisons of genetic variants with associated phenotypic findings suggest that mutations that affect nucleotide or microtubule binding strongly correlate with clinical severity (Boyle et al., 2021). However, mutations in the C-terminal cargo binding domains can also be pathogenic. Further, it can be challenging to draw structure/function conclusions based on variant position alone. Within a five amino acid stretch in the motor domain of KIF1A, variant p.V8M leads to hyperactive motility, p.R11Q causes hypoactive motility, and p.R13H inhibits microtubule binding (Boyle et al., 2021; Chiba et al., 2019). These findings highlight the importance of functional characterization of pathogenic *KIF1A* variants.

The three alleles tested in this study are predicted to differentially alter function. Analysis of the null allele (p.C92*) allows us to query the cellular effects of KIF1A loss in human neurons. The hypoactive p.P305L allele investigated here is known to impair KIF1A motor function (Kaur et al., 2020). This mutation localizes to a conserved motif adjacent to the microtubule-binding K-loop of KIF1A. Consistent with this, mutation of this residue inhibits microtubule binding, reducing motor velocity by ∼50% and decreasing motor run lengths by ∼25% in single molecule assays (Lam et al., 2021). In contrast, introduction of the p.R350G mutation led to a significantly higher landing rate for the binding of KIF1A to microtubules and a 3-fold faster velocity than wild type KIF1A (Chiba et al., 2019). These observations suggest the p.R350G mutation disrupts auto-inhibition of KIF1A as well as activating motility (Chiba et al., 2019).

### Effects of KIF1A mutations on neuron viability and neurite outgrowth *in vitro*

We tested the effects of these three mutations in iPSC-derived human neurons, focusing on homozygous lines as previous studies have found that the behavior of homodimeric mutants is a better predictor of clinical severity than the analysis of heterodimeric motors (Rao et al., 2025). Furthermore, hyperactivating mutants, such as p.R350G, are inherited as recessive mutations. Consequently, the homozygous condition represents the pathogenic state. KIF1A has previously been implicated in cell viability, with neuronal degeneration and death observed in KIF1A knockout mice both *in vivo* and *in vitro* primary neuronal cultures (Yonekawa et al., 1998). In our human iNeuron system, however, only hypoactive p.P305L mutants exhibited increased cell death as determined by CellEvent Caspase-3/7 detection assay, with no change in viability observed for null p.C92* or hyperactive p.R350G iNeurons. In primary cultures, Yonekawa *et al*. found that low level glutamate exposure could rescue neuronal death caused by KIF1A deletion, presumably by overcoming insufficient stimulation. Differences in culture conditions may explain why human iNeurons and mouse primary neurons lacking KIF1A exhibit different viability outcomes. Specifically, the BrainPhys medium used for iNeuron culture is designed to support synaptic development (Bardy et al., 2015), and may provide an environment supportive for neuronal viability compared to Neurobasal medium. These data reveal that under these conditions, human iNeurons can compensate for *KIF1A* loss to prevent premature cell death. It is interesting, therefore, that even in this permissive environment, hypoactive *KIF1A-*p.P305L iNeurons still exhibit an increase in cell death. This suggests that the p.P305L variant acts as a dominant negative in terms of neuron viability, with hypoactive KIF1A function diminishing a neuron’s chance of survival.

Somewhat surprisingly, delayed neurite outgrowth was observed in the two loss-of-function lines (p.C92* and p.P305L) but not in the p.R350G line. Recent work from the Bonifacino lab suggests a critical role for kinesin motors, both kinesin-1 and KIF1A, in the trafficking of lysosomes and associated RNA granules along the axon (De Pace et al., 2024). It is possible that the delay in neurite outgrowth seen in the loss-of-function lines may reflect an initial failure to correctly localize either lysosomes or mRNAs during early neurite outgrowth in KIF1A-null neurons. Since we observed a delay and not a block in outgrowth, kinesin-1 may compensate for KIF1A function at later stages of neurite extension.

An additional unexpected observation was the differences observed in KIF1A expression as measured by either western blot or qPCR. Immunoblot analysis indicates that the p.C92* missense mutation does equate to a KIF1A null line when homozygous, as predicted. The low levels of KIF1A mRNA detected by qPCR are consistent with nonsense-mediated decay. In contrast, we saw wild type-levels of KIF1A protein expression in the p.P305L and p.R350G lines, but significantly lower levels of KIF1A mRNA expressed in either line, relative to wild type. As KIF1A protein has been shown to undergo uniquely rapid degradation compared to other kinesin motors (Huang et al., 2020), we can speculate that the mutant motors may have a longer half-life, potentially due to either prolonged cargo interactions (p.P305L) or distal accumulation in neurite tips for the hyperactive motor (p.R350G). If there is indeed enhanced stability of the mutant motor relatively to wild type KIF1A in heterozygous patients, this would result in a more strongly dominant-negative phenotype, perhaps explaining the observations of Gennerich and colleagues that the biophysical properties of homodimeric mutants are more predictive of clinical severity than those of heterodimeric motors (Rao et al., 2025).

### Effects of KIF1A mutations on SVP trafficking

KIF1A has a critical function in the trafficking of synaptic vesicle precursors, conserved from *C. elegans* to mammalian neurons (Hall and Hedgecock, 1991; Hummel and Hoogenraad, 2021; Otsuka et al., 1991; Yonekawa et al., 1998). To test the effects of KIF1A mutations when expressed at endogenous levels in human neurons, we examined SVP trafficking using mScarlet-SYP (Guedes-Dias et al., 2019) via live-cell imaging. We found robust anterograde trafficking of SVPs in all genotypes examined. The absence of a significant deficit in anterograde velocity in p.C92* neurons, essentially null for KIF1A expression, is consistent with previous studies in mice indicating partial compensation by other kinesin superfamily members, most likely KIF1B*β* (Niwa et al., 2008). There is a trend toward slower anterograde velocities in the p.P305L neurons, consistent with a small dominant-negative influence on SVP trafficking.

More strikingly, there is a significant increase in the velocity of SVPs in p.R350G neurons, fully consistent with the identification of this mutation as a hyperactive allele. Quite unexpectedly, the R350G mutation in KIF1A also induced a significant increase in the retrograde velocity of SVPs along the axon. We also compared vesicle flux in both the anterograde and retrograde directions. As with velocity measurements, loss of KIF1A in the p.C92* neurons had no effect on SVP flux either outward or inward along the axon, but expression of the p.P305L mutation was sufficient to significantly inhibit both anterograde and retrograde flux. In contrast, the hyperactivating p.R350G mutation significantly increased flux in both the anterograde and retrograde directions. The ratio of retrograde to anterograde transport remained consistent across phenotypes, suggesting that SVP transport is carefully balanced along the axon, by mechanisms that are not yet understood. Finally, we compared the fluorescent intensity of motile SVP puncta, and found significant differences for both the C92* and P305L lines. These data suggest that while KIF1B*β* can partially compensate for loss of KIF1A function, KIF1B*β* may not be as effective in generating SVP transport carriers via sorting at the trans-Golgi network (Park et al., 2023). Unique kinetic properties of KIF1A motors, specifically their ability to remain engaged with the microtubule under hindering loads (Pyrpassopoulos et al., 2023), may also contribute to the increase in SVP intensity when functional KIF1A is present.

These findings of paired changes to anterograde and retrograde transport echo longstanding observations on the close coupling of anterograde and retrograde motor activity in neurons (Barkus et al., 2008; Brady et al., 1990; Pilling et al., 2006; Waterman-Storer et al., 1997). *In vitro* studies using reconstituted proteins demonstrate that KIF1A is an exclusively plus-end directed motor and have not predicted inhibitory effects on retrograde transport velocities. It is likely these minimal systems do not fully replicate the complex regulatory mechanisms controlling cargo movement in cells. However, in model organisms, some bi-directional effects on transport have been noted. In *C. elegans*, KIF1A mutations that activate anterograde flux inhibit retrograde flux (Chiba et al., 2019) but in Drosophila, loss of KIF1A induced decreased velocities and run lengths for retrograde-moving dense core vesicles, although not for synaptotagmin-positive organelles (Barkus et al., 2007). More detailed studies will be required to fully interrogate the mechanisms involved to see if they involve biophysical aspects such as changes in the assembly of motor teams or changes in drag forces from opposing motors, or instead reflect changes in the mechanisms that control directional switching of SVP trafficking in neurons.

### Localized delivery of synaptic components is differentially affected by mutations in KIF1A

Elegant *in vivo* studies have shown that loss of KIF1A leads to a depletion of synaptic vesicles from presynaptic sites, first in *C. elegans* (Hall and Hedgecock, 1991; Otsuka et al., 1991) and then in mouse (Yonekawa et al., 1998). Fully consistent with these studies, we find that either loss of KIF1A (p.C92* mutant) or expression of the p.P305L missense mutation in KIF1A leads to the marked accumulation of presynaptic markers in the soma of human neurons, and a depletion of these markers from heterologous synapses. In striking contrast, the hyperactive R350G mutant effectively depletes presynaptic markers from the soma and enriches these markers at heterologous presynapses.

Interestingly, while loss-of-function mutations p.C92* and p.P305L often converge on shared cellular outcomes, such as somal retention of presynaptic cargo and decreased synapse density, they arrive at these outcomes through distinct pathways. While p.P305L hypoactive motors induce a predicted reduction in axonal SVP flux and velocity, p.C92* axons lacking KIF1A motors show no deficits in axonal trafficking parameters but instead exhibit reduced intensity of motile synaptic cargos. However, both show a similar downstream effect on trafficking to presynaptic sites, in stark contrast to the hyperactive R350G allele.

Recent studies have identified a key role for localized microtubule dynamics in the delivery and exchange of SVPs within presynaptic sites such as *en passant* synapses (Aiken and Holzbaur, 2024; Guedes-Dias et al., 2019; Qu et al., 2019). While axonal microtubules are classically considered to form stable arrays in the axon (Baas et al., 1988), live imaging of microtubule dynamics has shown that there are localized bursts of plus-end polymerization of microtubules at sites coincident with the stable accumulation of presynaptic markers such as synaptophysin and synaptobrevin (Guedes-Dias et al., 2019). Motile SVPs preferentially pause and are retained at these sites, while perturbation of microtubule dynamics is sufficient to disrupt further SVP accumulation (Guedes-Dias et al., 2019; Aiken and Holzbaur, 2024). As KIF1A is a highly processive motor both *in vitro* and *in vivo* (Benoit et al., 2024; Guedes-Dias et al., 2019; Lee et al., 2003; Pyrpassopoulos et al., 2023; Soppina et al., 2014), current models predict that this motor will traverse to the end of the ‘track’—the plus end of the microtubule—leading to preferential detachment and cargo delivery at microtubule plus end-enriched presynaptic sites (Aiken and Holzbaur, 2024). We asked how this mechanism might be affected by mutations in KIF1A by monitoring both microtubule dynamics and SVP delivery in wild-type or KIF1A mutant neurons. Across the genotypes tested, we saw similar densities of microtubule dynamic hotspots. In both p.C92* and p.P305L neurons, we saw similar correlations between these microtubule hotspots and sites of local SVP accumulation. In striking contrast, however, we found this close correlation was lost in neurons expressing the hyperactive p.R350G mutation. This finding is supported by the significantly higher microtubule landing rate (approximately 10-fold increase compared to wild-type motor) previously reported for p.R350G motors (Chiba et al., 2019), allowing hyperactive KIF1A motors to rapidly bind neighboring microtubules following plus end dissociation. Based on these data, we hypothesize that the p.R350G mutation disrupts the ability of KIF1A to read and respond to the local microtubule landscape characteristic of presynaptic sites.

### KIF1A loss-of-function vs. gain-of-function mutants lead to discrete functional changes in the timing of synaptic development

Human neurons in culture form functional synapses over time. While slower and less efficient than the synapse formation observed for primary rodent neurons, iPSC-derived human neurons exhibit significant apposition of pre- and post-synaptic markers by DIV42, a process that is enhanced by co-culture with glial cells (Aiken and Holzbaur, 2024). We examined synapse formation in *KIF1A* mutant neurons, and found that all three mutant lines tested were less effective than control neurons in balancing apposition of pre- and post-synaptic compartments. This led us to query synaptic function across these lines using MEA technology, and found clear deficits in mean firing rates, burst activity, and network burst activity for neurons in co-culture with glia. While loss-of-function mutants p.C92* and p.P305L led to a maturation defect in synaptic connectivity, the gain-of-function hyperactive p.R350G mutant induced an acceleration of synaptic activity, with precocious network bursts recorded at early stages. This contradictory impact on the timing of functional synaptic connections aligns with the differential effects observed for SVP trafficking and local SVP accumulation at presynaptic sites enriched in microtubule plus ends. These data suggest that the density of stable axonal SVP+ sites does not impact spontaneous synaptic vesicle release, as *KIF1A* mutant axons that contain appropriate SVP+ site density (p.C92* and p.P305L) exhibit decreased mean firing rates. Further, axons of neurons expressing the hyperactive p.R350G mutant display increased motile SVP cargos but fewer SVP+ sites. These changes do not dramatically impact individual spike or burst frequencies, but enhance overall network connectivity as revealed by precocious network bursts. It is tempting to speculate that this change in network wiring may be a consequence of presynaptic size/strength at axon terminals. A potential buildup of presynaptic material in distal regions by the hyperactive p.R350G motors may accommodate larger axonal terminals, resulting in enhanced post-synaptic potential and increased likelihood of synchronized, network-wide bursts of activity (Lavi et al., 2015). This acceleration in network formation comes at a cost, however, as these cultures experiencing a sharp decline in neuronal activity over time, potentially due to excitotoxic effects.

### Conclusions

Technological developments in gene-editing and differentiation of human iPSC-derived neurons now allow detailed phenotypic testing of pathogenic human mutations affecting neuronal function. Here, we directly compared the effects of three disease-causing mutations on neuronal viability, neurite outgrowth, microtubule dynamics, SVP trafficking, synapse formation, and synaptic function. While each mutation induced a distinct subset of molecular and cellular changes, the two loss-of-function variants, p.C92* and p.P305L, converged on a shared functional deficit in synaptic maturation. This convergence may offer hope for therapeutic strategy tailored to specific function changes to KIF1A motor properties. As there are nearly 200 known *de novo* pathogenic mutations identified to date within the *KIF1A* gene, the potential for shared treatments is of high interest although our findings instead suggest that individually-targeted therapeutic interventions are more likely to be successful. While we focused on detailed analysis of homozygous mutant lines, it will be of interest to more fully test heterozygous lines modeling de novo mutations, as well as to expand testing to other pathogenic variants. We anticipate these studies may identify convergence on downstream consequences such as synaptic dysfunction, and provide further insights into therapeutic interventions to ameliorate functional deficits caused by specific *KIF1A* variants.

### Author Contributions

J.A., C.B. and E.L.F.H. conceived the project and designed experiments. J.A., C.B., and N.M. performed the experiments and analyzed the data, and J.A. and E.L.F.H wrote and edited the manuscript with contributions from C.B., N.M., and B.L.P.

## Acknowledgements

The authors gratefully acknowledge support from NIH awards to J.A. (NINDS F32NS117672) and E.L.F.H. (NIGMS R35 GM126950).

## Declaration of Interests

The authors declare no competing interests.

## RESOURCE AVAILABILITY

### Lead contact

Further information and requests for resources and reagents should be directed to and will be fulfilled by the lead contact, Erika Holzbaur.

### Materials availability

The gene-edited cell lines used here are available from JAX and the plasmids generated in this study are available upon request.

### Data and code availability

Excel datasets for all graphs have been uploaded to Zenodo, doi: 10.5281/zenodo.16924227. Any additional information required to reanalyze the data reported in this work paper is available from the Lead Contact upon request.

## EXPERIMENTAL MODEL AND SUBJECT DETAILS

### Human induced pluripotent stem cells (iPSCs)

KOLF2.1J-background wild-type and *KIF1A*-p.C92*, -p.P205L, and -p.R350G knock-in iPSCs were a gift from the Cellular Engineering Team led by Justin McDonough (The Jackson Laboratory for Genomic Medicine, Farmington, CT, USA) as part of the KIF1A.org iPSC resource initiative. The KOLF2.1J parental line has been characterized previously (Pantazis et al., 2022). iPSCs were maintained on Growth Factor Reduced Matrigel-coated plates (Corning) and cultured in Essential 8 medium (Thermo Fisher Scientific), changed daily. To generate stable iPSC lines expressing doxycycline-inducible human NGN2 (hNGN2) via a PiggyBac transposon system, cells were transfected with the PB-TO-hNGN2 plasmid (a gift from M. Ward, NIH, MD, USA) and transposase at a 1:2 ratio (transposase:vector) using Lipofectamine Stem (Thermo Fisher Scientific). After 72 hours, transfected cells were selected with 0.5 μg/ml puromycin (Takara) for 48 hours. Differentiation into induced neurons (iNeurons) was performed following established protocols (Fernandopulle et al., 2018; Pantazis et al., 2022). Briefly, iPSCs were passaged using Accutase (Sigma-Aldrich) and replated on Matrigel-coated dishes in Induction Media, consisting of DMEM/F12 supplemented with 1% N2 supplement (Gibco), 1% non-essential amino acids (NEAA; Gibco), and 1% GlutaMAX (Gibco), with 2 μg/ml doxycycline. After 72 hours of doxycycline treatment, iNeurons were dissociated with Accutase and cryopreserved in liquid nitrogen. The full protocol is available on Protocols.io: https://doi.org/10.17504/protocols.io.261ge348yl47/v1.

It is worth noting that the KOLF2.1J iPSC line has been reported to have only one functional allele of neuronally expressed genes *JARID2* and *ASTN2* (Gracia-Diaz et al., 2024; Ryan et al., 2024). While chromosomal deletions encompassing these genes have been linked to neurodevelopmental disorders, mice heterozygous for a deletion encompassing *Jarid2* or *Astn2* in the absence of other gene deletions exhibit no obvious neurodevelopmental phenotypes (Chang et al., 2015; Zhang et al., 2023). The widely distributed and well-characterized KOLF2.1J iPSC line therefore remains a useful tool for the neurodevelopmental research community.

### Culture and transfection of iPSC-derived neurons in monoculture

Cryopreserved, pre-differentiated iNeurons were thawed and plated onto 35-mm glass-bottom imaging dishes (MatTek; P35G-1.5-20-C) pre-coated with 100 µg/mL poly-L-ornithine (Sigma-Aldrich; P3655), at a density of 3x10⁵ cells per dish. For each experimental condition, three to four independent cultures were prepared using at least two different neural induction batches. iPSC-derived neurons were maintained in iNeuron Maintenance Media composed of BrainPhys™ Neuronal Medium (STEMCELL Technologies; 05790), supplemented with 2% B27 Supplement (Gibco™/Thermo Fisher Scientific; 17504-044), 10 ng/mL BDNF (PeproTech; 450-02), 10 ng/mL GDNF (PeproTech; 450-10), 10 ng/mL NT-3 (PeproTech; 450-03), and 1 mg/mL Mouse Laminin (Thermo Fisher Scientific; 23017-015). To prevent non-neuronal cell growth, iNeuron Maintenance Media was initially supplemented with 10 µM 5-Fluoro-2’-deoxyuridine (FUdR; Sigma Aldrich; F0503). Media was partially refreshed weekly by replacing 40% with fresh, pre-equilibrated iNeuron Maintenance Media. Unless specified otherwise, live imaging and immunocytochemistry were carried out 21 days post-thaw (DIV21). For live-cell imaging, iNeurons were transfected using Lipofectamine Stem Transfection Reagent (Thermo Fisher Scientific; STEM00001) and 0.75–2 µg of total plasmid DNA. A detailed protocol for culturing and transfecting iPSC-derived neurons for live imaging is available on Protocols.io: https://doi.org/10.17504/protocols.io.x54v9dj4zg3e/v1.

### Rat primary astrocyte isolation and culture

All procedures were conducted in accordance with the guidelines of the Institutional Animal Care and Use Committees at The Children’s Hospital of Philadelphia and the University of Pennsylvania. Primary rat astrocytes were isolated from postnatal day 1 Sprague Dawley rat pups (Charles River Laboratories, Wilmington, MA; RRID: RGD_737891) and plated in poly-D-lysine-coated T75 flasks. Cells were initially cultured in Neurobasal medium (Gibco™/Thermo Fisher Scientific; 21103049) supplemented with 2% B27 at 37°C in a 5% CO₂ incubator, following previously described methods (Jensen et al., 2015). After 24 hours, the medium was replaced with Neurobasal medium containing B27, 10 ng/mL basic fibroblast growth factor (bFGF), 2 ng/mL platelet-derived growth factor-AA (PDGF-AA), and 1 ng/mL neurotrophin-3 (NT-3). Upon reaching confluence (around DIV7), astrocytes were separated from oligodendroglial cells using the established “shake-off” method (McCarthy and de Vellis, 1980). Remaining astrocytes were passaged into fresh poly-D-lysine-coated T75 flasks using Trypsin-EDTA (Gibco™/Thermo Fisher Scientific; 25300-054) and cultured in astrocyte growth medium consisting of DMEM (Corning; 10-017-CM) supplemented with 10% fetal bovine serum (FBS; HyClone; SH30071.03), 1% non-essential amino acids (NEAA), and 1% Penicillin-Streptomycin (Pen/Strep). Media was refreshed weekly until cells reached full confluence, typically by DIV14. At this point, primary astrocytes were cryopreserved in astrocyte growth medium supplemented with 10% dimethyl sulfoxide (DMSO).

### Co-culture of iPSC-derived neurons with primary rat astrocytes

Primary rat astrocytes were thawed into uncoated T75 flasks and maintained in astrocyte growth medium at 37°C in a 5% CO₂ incubator. Upon reaching confluence (approximately 7 days post-thaw), cells were passaged using Trypsin-EDTA and seeded onto glass coverslips within 6-well plates at a density of 50,000 cells per well. Once astrocytes reached confluence on the coverslip (typically 4–7 days post-passage), 100,000 cryopreserved, pre-differentiated iNeurons were thawed and plated onto the astrocyte monolayer in pre-equilibrated Neuron Maintenance Media. Media was partially refreshed weekly by replacing 40% with fresh, pre-equilibrated Neuron Maintenance Media. Immunocytochemistry was performed at 42 days after iNeuron thawing (corresponding to DIV42).

### Human embryonic kidney cells

Human embryonic kidney 293T cells (HEK293T; referred to as HEK cells) were authenticated by short tandem repeat (STR) profiling. Cells were maintained in DMEM (Corning; 10-017-CM) supplemented with 10% fetal bovine serum (FBS) and cultured on standard 10-cm sterile tissue culture plates at 37°C in a humidified, 5% CO₂ water-jacketed incubator. Cells were passaged using trypsin once they reached 60–90% confluency.

## METHOD DETAILS

### Neuron viability assessment

CellEvent™ Caspase-3/7 Green Detection Reagent (Thermo Fisher Scientific; C10723) was added to DIV7 iNeurons at a 1:400 dilution in Imaging Media consisting of low fluorescence Hibernate A Medium (Brain Bits®; HA100) supplemented with 2% B27, 10 ng/mL BDNF, 10 ng/mL NT-3, and 1 mg/mL Mouse Laminin, and incubated for 20 minutes at 37°C. Immediately following incubation, CellEvent™ dye was removed and cells were stained with Hoechst 33342 (Thermo Fisher Scientific) diluted 1:20,000 in Imaging Media. Brightfield, 405 nm, and 488 nm fluorescence images were acquired using a Leica inverted fluorescence microscope equipped with appropriate excitation/emission filter sets and a HC PL APO 20×/0.70 dry objective. Images were analyzed using ImageJ to quantify the percentage of cells with activated caspase-3/7 (indicated by bright green nuclei) and total cells (indicated by Hoechst+ nuclei).

### Live imaging of iNeurons

iNeurons plated on 35-mm glass-bottom imaging dishes were imaged in an environmental chamber maintained at 37°C using iNeuron Imaging Media (described above). Time-lapse imaging data were acquired using a Hamamatsu ORCA-Fusion C14440-20UP camera operated via VisiView software. Axonal imaging regions were selected based on morphological criteria—specifically, long, uniformly thin processes with no visible swellings, located at least ∼200 μm from the soma. For synaptic vesicle precursor (SVP) motility experiments, one photobleaching cycle was performed with the 405 nm laser for 3 ms/pixel using a ViRTEx Realtime Experiment Control Device before collecting single-channel time-lapse images of SVPs (mScarlet-SYP) at a frame rate of 200 ms per frame for 5 minutes. For imaging microtubule plus-end dynamics, two-channel acquisition of MACF43-GFP and mScarlet-SYP was performed at 1-second intervals for 5 minutes. SVP+ site location and size were determined using kymograph analysis from the mScarlet-Syp or mScarlet-Syb channel, where SVP+ sites were defined as stationary (vertical) signals that remained consistently above background for the entire duration of the kymograph.

### RNA Extraction and Quantitative Real-Time PCR (qRT-PCR)

Total RNA was extracted using TRIzol reagent (Invitrogen; 15596026) following the manufacturer’s guidelines with minor modifications. Briefly, cell culture media was aspirated, and 150 μL of TRIzol was added directly to adherent DIV21 iNeurons, grown at a density of 800K per well of a 6-well plate. For phase separation, samples were incubated at room temperature for 5 minutes, followed by the addition of 30 μL chloroform and 2 μL GlycoBlue (Thermo Fisher; AM9515) for visualization. Tubes were vortexed for 10 seconds, incubated on ice for 10 minutes, and centrifuged at 12,000×g for 15 minutes at 4 °C. The resulting upper aqueous phase, containing RNA, was carefully transferred to a new tube. RNA was precipitated by adding 75 μL isopropanol to the aqueous phase, inverting to mix, and incubating for 10 minutes at room temperature. Samples were centrifuged at 12,000×g for 10 minutes at 4 °C. The RNA pellet was washed by resuspension in 45 μL of 75% ethanol, followed by brief vortexing and centrifugation at 7,500×g for 5 minutes at 4 °C. The supernatant was discarded, and the pellet was air-dried for 5–10 minutes until it began to clarify but was not completely dry.

Quantitative reverse transcription PCR (qRT-PCR) was performed using the Brilliant II SYBR® Green QRT-PCR 1-Step Master Mix Kit (Agilent Technologies; NC9630749) according to the manufacturer’s instructions. Each 25 μL reaction contained the following components: 12.5 μL of 2× SYBR Green QRT-PCR Master Mix, 0.375 μL of 100× RT/RNase block enzyme mixture, 0.3 μM of each forward and reverse primer, and 50 ng of template RNA, with RNase-free water added to adjust the final volume. Reactions were assembled on ice and gently mixed before being loaded into a real-time PCR instrument. Thermal cycling was performed with the following conditions: reverse transcription at 50 °C for 30 minutes, initial denaturation at 95 °C for 10 minutes, followed by 40 cycles of 95 °C for 30 seconds and 60 °C for 1 minute. A dissociation curve analysis was performed at the end of each run to confirm amplification specificity. Non-template controls (NTC) were included to monitor for contamination. Relative gene expression was calculated using the ΔΔCt method, with normalization to GAPDH housekeeping gene expression.

### Immunocytochemistry of iNeurons

For immunocytochemistry of neurite outgrowth, endogenous SVPs, synapses, or heterologous synapses, iNeurons cultured on 35-mm glass-bottom imaging dishes or glass coverslips were fixed with 4% paraformaldehyde/4% sucrose in PBS for 10 minutes at room temperature. Following three PBS washes, cells were permeabilized and blocked for 1 hour at room temperature in Blocking Solution containing 1% bovine serum albumin (BSA), 5% goat serum, and 0.1% Triton X-100 in PBS. Neurons were then incubated overnight at 4°C with primary antibodies diluted in Blocking Solution. The following primary antibodies were used: NF-H (mouse anti-NF-H; BioLegend, 801601; 1:1000), Synapsin I/II (guinea pig anti-Synapsin I/II; Synaptic Systems, 106-004; 1:1000), PSD-95 (rabbit anti-PSD-95; Synaptic Systems, 124-008; 1:500), MAP2 (mouse anti-MAP2; EMD Millipore, MAB3418; 1:200), Synaptophysin (mouse anti-Synaptophysin; Sigma-Aldrich, S5768; 1:200), and Synaptobrevin-2 (rabbit anti-Synaptobrevin-2; Cell Signaling, 13508; 1:250). After three additional PBS washes, cells were incubated for 1 hour at room temperature, protected from light, with species-specific secondary antibodies (1:500 dilution) prepared in Blocking Solution. Following a final set of three PBS washes, coverslips were mounted using ProLong Gold Antifade Mountant (Thermo Fisher; P36930). Images were acquired as z-stacks with a 200 nm step size using the Hamamatsu ORCA-Fusion C14440-20UP camera operated via VisiView software, as described above.

### Heterologous synapse assay

HEK cells transfected with pBI-BFP-NL1 (bi-cistronic vector expressing untagged NL1 and cytosolic BFP) using 6 µl FUGENE:1 µg DNA and were added to DIV13 iNeurons cultured in 35-mm imaging dishes 24 hours after transfection (100K transfected HEK cells added to 300K DIV13 iNeurons). 24 hours after the addition of the transfected HEK cells, at iNeuron DIV14, cells were fixed for immunocytochemistry in 4% paraformaldehyde supplemented with 4% sucrose and stained as described above.

### Immunoblotting

iNeuron samples were lysed in RIPA buffer (50 mM Tris-HCl, 150 mM NaCl, 0.1% Triton X-100, 0.5% deoxycholate, 0.1% SDS) supplemented with 2× Halt Protease and Phosphatase Inhibitor Cocktail. Lysates were centrifuged at 18,000 × g for 15 minutes to remove cellular debris, and the protein concentration of the supernatant was quantified using the Pierce™ BCA Protein Assay Kit (Thermo Fisher Scientific; 23225). Proteins were resolved on Mini-PROTEAN TGX Stain-Free Gels (4-15% gradient; Bio-Rad: 4568084) gels and transferred to Immobilon®-FL PVDF membranes (Millipore; 05317) using a wet transfer system (Bio-Rad). Membranes were stained for total protein using LI-COR Revert 700 Total Protein Stain and imaged with the Odyssey CLx Infrared Imaging System (LI-COR). After imaging, membranes were de-stained and blocked for 5 minutes with EveryBlot Blocking Buffer (Bio-Rad; 12010020). Primary antibody incubations were carried out overnight at 4°C in EveryBlot Blocking Buffer using the following antibodies: anti-KIF1A (mouse; BD Biosciences, 612094; 1:200), anti-synaptophysin (guinea pig; Synaptic Systems, 101004; 1:1000), and anti-synaptobrevin-2 (rabbit; Cell Signaling, 12508; 1:1000). Membranes were then washed three times with TBS containing 0.1% Tween-20 (TBST) and incubated for 1 hour at room temperature with species-appropriate secondary antibodies diluted 1:20,000 in EveryBlot with 0.01% SDS. After three additional washes with TBST and one final wash with TBS, membranes were imaged using the Odyssey CLx system. Western blot data were analyzed using LI-COR Image Studio Software, and protein levels were quantified relative to total protein and compared between wild type and *KIF1A* mutant iNeurons. A detailed protocol is available on Protocols.io: https://doi.org/10.17504/protocols.io.5jyl8j5zrg2w/v1. Data represent three independent experiments (N = 3), each from separate iNeuron induction batches.

### Multi-electrode array (MEA) recordings of spontaneous synaptic activity

Spontaneous neuronal activity was recorded from iNeuron monocultures and iNeuron/primary rat astrocyte co-cultures using the Multiwell-MEA-System, Multiwell-MEA workstation with temperature control from Harvard Bioscience, Multichannel Systems. 24-well plates with PEDOT electrodes on glass (Harvard Biosciences; 24W300/30G-288) were coated with 100 µg/mL poly-L-ornithine (Sigma-Aldrich; P3655) overnight at 4°C, followed by 20 µg/ml Mouse Laminin (Thermo Fisher Scientific; 23017-015) for 1 hour at 37°C prior to cell plating. For iNeuron monocultures, 150K iNeurons were seeded per well. For iNeuron/glia co-cultures, primary astrocytes where seeded onto the precoated electrode plates (20K per well) and allowed to reach confluency, approximately 5 days. iNeurons were thawed and plated on top of the primary astrocytes (50K per well). Longitudinal recordings were performed starting at DIV7 using Harvard Biosciences MEA recording system (Multiwell-MEA-System) and recorded every other day or two until DIV44. Cultures were maintained in Neuron Maintenance Media, with medium partially refreshed following MEA recording by replacing 40% with fresh, pre-equilibrated Neuron Maintenance Media. For data acquisition, MEA plates were equilibrated in the Multiwell-MEA-System recording chamber maintained at 37°C for 10 minutes. Signals from 12 electrodes per well were recorded for 10 minutes simultaneously at a sampling rate of 20 kHz per channel using Multiwell-Screen software (Multi Channel Systems) under spontaneous conditions without pharmacological treatments.

## QUANTIFICATION AND STATISTICAL ANALYSIS

### Kymograph generation and tracing

Kymographs from live-imaging time series were generated using the KymographClear plugin in ImageJ (Schindelin et al., 2012), following published guidelines (Mangeol et al., 2016), with the line width set to 3 pixels. Axonal segments approximately 100 µm in length that exhibited active synaptic vesicle precursor (SVP) trafficking and/or microtubule comet activity were selected for analysis. Within each kymograph, individual motile tracks were manually traced in ImageJ, and the coordinates of each inflection point were recorded in an Excel spreadsheet. Pixel coordinates were then converted to micrometers (x-axis) and seconds (y-axis) for quantitative analysis. The resulting distance (x) and duration (y) values were used to calculate parameters of SVP transport. SVP+ site positions and lengths were determined based on the locations of bright, stationary mScarlet-Syb puncta along the axon.

### SVP movement analysis

Motile SVPs were classified as anterograde if they exhibited a net displacement greater than 10 µm in the anterograde direction, or retrograde if their net displacement exceeded 10 µm in the retrograde direction. SVP flux (vesicles per minute) was calculated by counting the number of anterograde or retrograde SVPs entering the kymograph and normalizing to the total imaging duration. The coordinates of SVP tracks were used to calculate instantaneous velocity, and the five fastest velocities were selected to determine “maximum velocity” per axon. N=3 experimental replicates, n≥12 axons and n≥74 motile SVPs. To determine motile SVP intensity, a maximum intensity measurement (with background subtracted) was recorded as soon an as an anterogradely moving SVP entered the imaging field.

### Microtubule comet analysis

Kymographs were analyzed for microtubule comet activity and SVP+ site density. SVP+ regions were determined by the position of stable (vertical) signal of Syb signal in mScarlet-Syp kymographs. Microtubule comet density (comets/µm/min) were determined by manually tracing the comet events within the kymograph and normalizing to the kymograph length and time. Microtubule comets associated with SVP+ sites were determined by counting the number of events that occur within 10 µm of the centroid of an SVP+ site. SVP+ sites with microtubule comet activity were quantified by counting the number of SVP+ sites that experienced at least one microtubule comet event starting, ending, or passing through.

### SynapseJ analysis

Z-stack images of iNeurons at DIV42, co-cultured with primary rat astrocytes, were immunostained for Synapsin I/II, PSD-95, and MAP2, and analyzed for synapse detection using SynapseJ (v.2) following the previously described protocol (Moreno Manrique et al., 2021). Detection thresholds for pre- and postsynaptic channels were set at two-thirds of the mean puncta intensity, as determined using ImageJ’s 3D Objects Counter. Default settings were used for noise and the Find Maxima function. Presynapse size was set to 0.2-2.5 µm, and postsynaptic size was set to 0.06-2 µm. After automated synapse identification by SynapseJ, the resulting merge.tif files were analyzed in ImageJ using the 3D Object Counter to quantify apposed pre- and postsynaptic puncta based on 3D masks generated by SynapseJ. Original z-stacks were reviewed to confirm that identified synaptic puncta were localized to MAP2-positive dendrites. MAP2-positive somatodendritic area was measured from thresholded maximum-projection images. Synapse density was calculated by dividing the number of matched pre/postsynaptic puncta by the MAP2-positive area per imaging field. The percentage of presynaptic puncta with a corresponding postsynaptic partner was calculated by dividing the number of “Synapse Pre” objects by the total “Pre No.” objects. Data represent 3 independent experiments, with at least 30 imaging fields analyzed per condition.

### Somatic SVP intensity and heterologous synapse intensity analysis

To measure the endogenous intensity of SVP proteins (synaptophysin and synaptobrevin-2) in the soma, the MAP2 channel was used to select an ROI around the somatic compartment. This ROI was used to measure the mean grey value of the somatic SYP signal using a sum projection. To determine intensity of these same SVP proteins within heterologous synapses formed within axonal regions crossing NL1+ HEK cells, ImageJ’s Thresholding tool was used to segment MAX-projection images and an 8-bit object mask based on the top 3% (for synaptophysin) or 2% (for synaptobrevin-2) intensity was generated. ImageJ’s Analyze Particles tool was used on the object mask redirected to the original image to determine intensity values. N=3 experimental replicates, n≥30 HEK cells encountering numerous crossing axons.

### MEA recording of spontaneous synaptic activity analysis

A high-pass filter (second order, 100 Hz), low pass filter (fourth order, 3500Hz), and a Notch Filter (50Hz) were applied to recorded data to isolate fast-spiking activity and remove high-frequency noise in the Multiwell Analyzer Software. Individual spikes were detected with a noise threshold of +/- 5 standard deviations from the baseline noise and exported. Exported spike detection files were analyzed in R Studio, utilizing in-house code to adapt files for compatibility with meaRtools, a published R package for MEA data analysis (available on request) to calculate each parameter as an average per well. Mean firing rates (spikes/second) were calculated by averaging firing rates from active electrodes, defined as electrodes exhibiting >10 spikes per minute. The following activity parameters were assessed: active electrodes, a readout of which of the 12 total electrodes experience at least 10 spikes/min, correlating to culture health and overall activity; mean firing rate, the average number of electrical spikes over the ten minute recording period; burst frequency and duration, defined as repeated spikes on an electrode that occur within 0.05s (50ms) and terminate after 0.1s (100ms) of inactivity; and network burst frequency and duration, defined as busts detected across at least 4 electrodes.

### SuperPlot data presentation and statistics

Experimental data were visualized using the SuperPlot format (Lord et al., 2020), with distinct colors representing the different genotypes and shapes representing individual biological replicates. Data points from each replicate (solid shapes) and their corresponding experimental means (larger, open shapes) were plotted using the shape with color set to 50% and 100% opacity, respectively. Figure legends specify the statistical tests used, and exact p-values are shown directly on the plots. For analyses involving linear mixed-effects (LME) models, R version 4.4.1 (2024-06-14) was used with the “nlme” package. Genotype was modeled as a fixed effect, while independent experiments or cultures were treated as random effects.

Unless otherwise noted in the figure legends, error bars represent standard deviation calculated from replicate means.

### Plasmids

Expression constructs used in this study include PB-TO-hNGN2 (Addgene; 172115), PiggyBac transposase vector, PGK mScarlet-Synaptophysin (Addgene; 206145), pCIG2-mScarlet-Syb-IRES-GFP-MACF43 (Aiken and Holzbaur, 2024), and pBI-NL1-BFP (Addgene; 206153). All constructs were verified by DNA sequencing.

---

**Figure S1.**
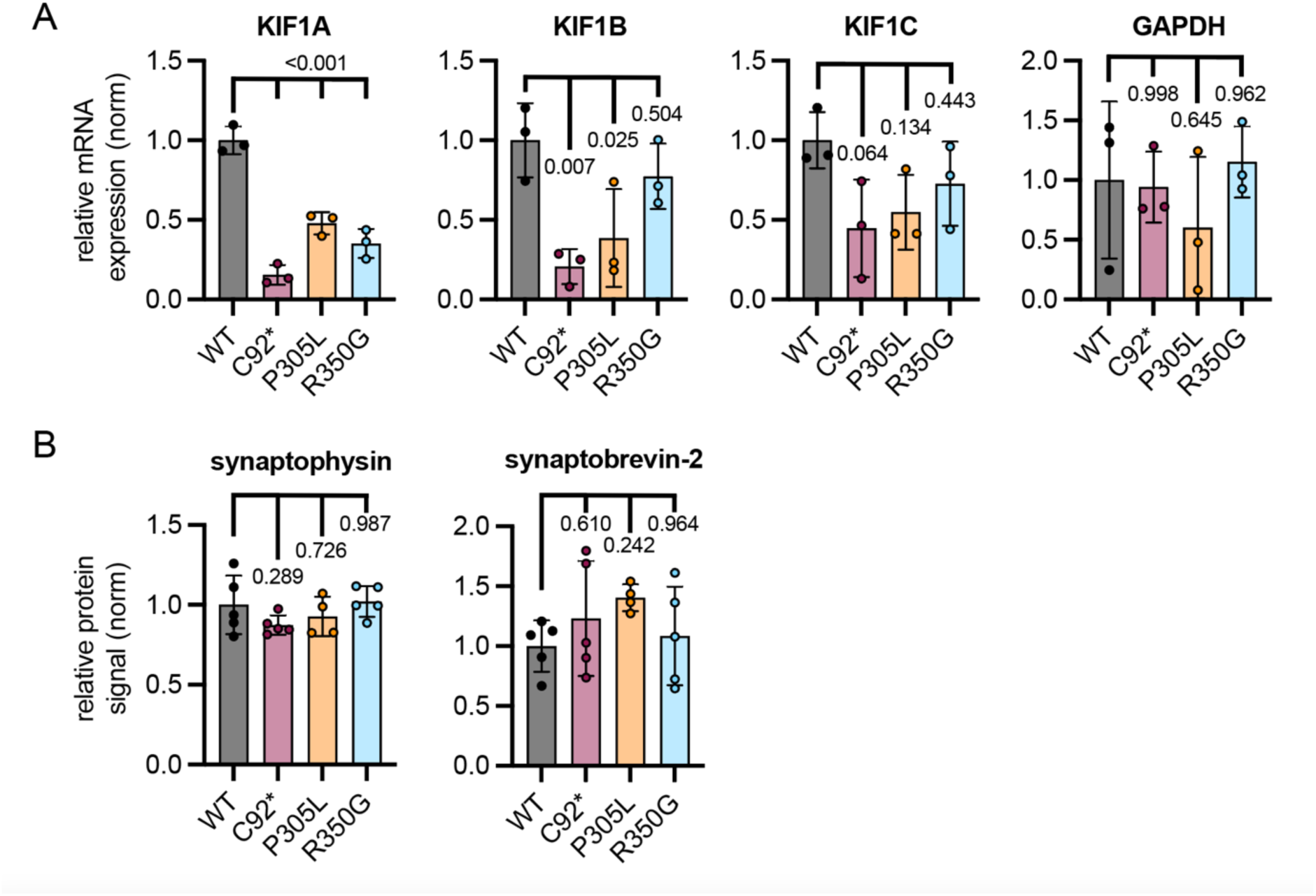
**(A)** RT-qPCR analysis reveals decrease in *KIF1A* mRNA level across *KIF1A* variant conditions. KIF1A null (p.C92*) also leads to decreased mRNA level of additional kinesin-3 motor family members, *KIF1B* and *KIF1C*. Housekeeping gene GAPDH was used as a reference control and wild-type condition was used to calculate relative fold change in kinesin-3 genes of interest. **(B)** Quantification of relative protein levels of synaptophysin (SYP) and synaptobrevin-2 (SYB2) in DIV21 iNeuron lysate, corresponding to example western blot displayed in Figure 3A (mean ± standard deviation; n = 4 independent experiments; reported *p*-values are from ANOVA with multiple comparisons).

**Figure S2.**
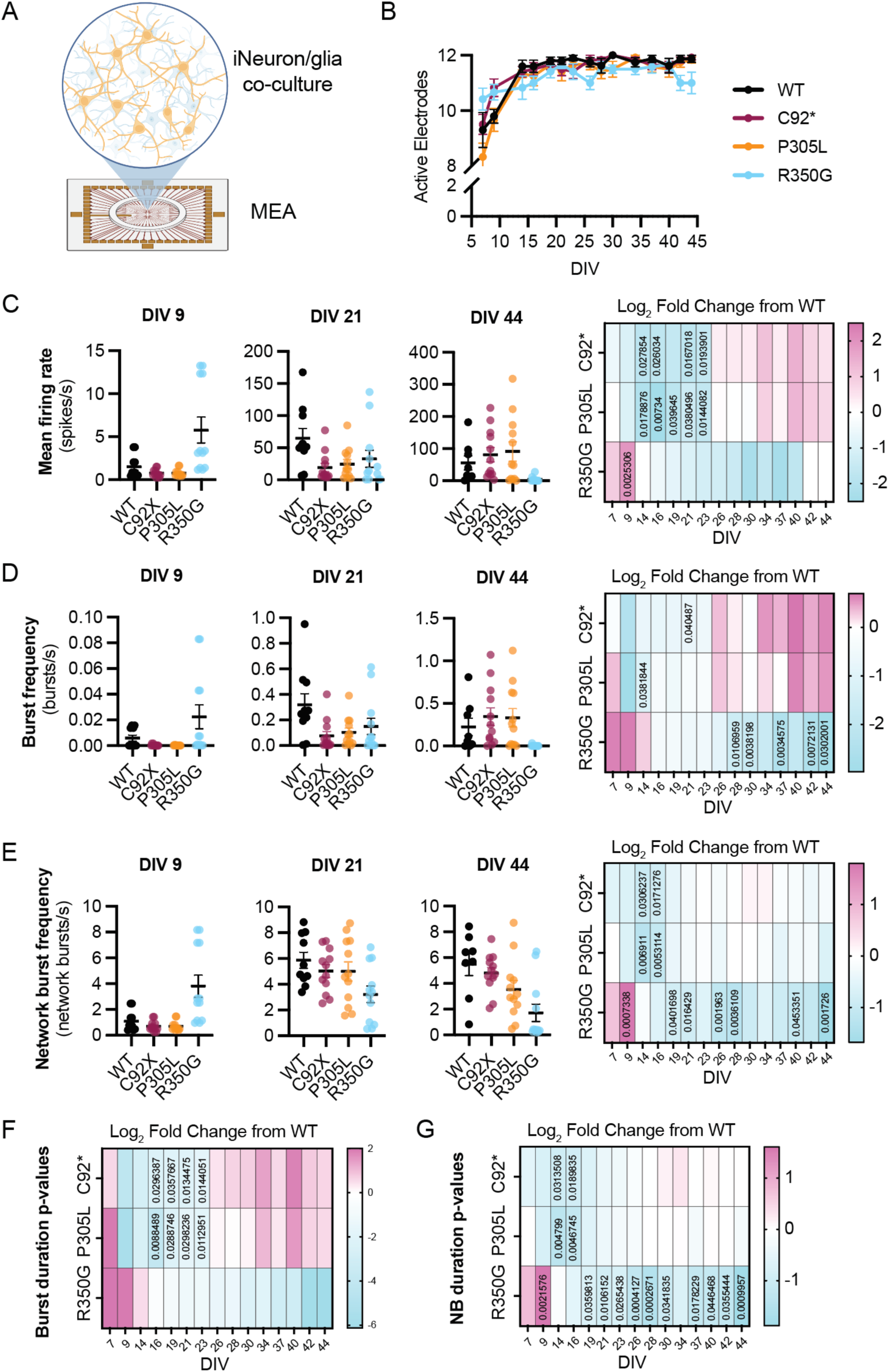
**(A)** Schematic of iNeuron/glia co-culture experimental set-up for spontaneous MEA recording. **(B)** Longitudinal tracking of active electrodes for wild-type KIF1A iNeurons and p.C92*, p.P305L, p.R350G mutant iNeurons. Hyperactive p.R350G iNeurons exhibit an increase in active electrodes compared to wild type at early stages of culture. **(C-E)** Mean firing rate (C), burst frequency (D) and network burst frequency (E) for KIF1A wild-type and mutant cultures at DIV 9, 21, and 44. Each dot represents an individual electrode, and error bars represent standard deviation. P-value heat map plots are provided to the right of individual time point plots. **(F, G)** P-value heat map plots for burst duration (F) and network burst (NB) duration (G). Heat map plots of p-values provide insight into significance across all recorded time points, with p-values < 0.05 provided in the corresponding box. Blue-shaded boxes reveal a decrease compared to wild type, and pink-shaded boxes reveal an increase compared to wild type.

## Notes

https://10.5281/zenodo.16924227

### Competing Interest Statement

The authors have declared no competing interest.

